# Multi-layered control of chromosomal assembly of the meiotic DNA break machinery

**DOI:** 10.64898/2026.06.21.733591

**Authors:** Masaru Ito, Stephen Mwaniki, Akira Shinohara, Kunihiro Ohta

## Abstract

Meiotic DNA double-strand breaks (DSBs) form in chromatin loops that are tethered to chromosome axes. How proteins required for DSB formation *in vivo* (DSB proteins) assemble on meiotic chromosomes to control non-random DSB distribution remains unsolved. Here we mapped the spatial distributions of all ten DSB proteins that form three complexes (Spo11-Ski8-Rec102-Rec104 [Spo11-core], Rec114-Mei4-Mer2 [RMM], and Mre11-Rad50-Xrs2 [MRX]) in budding yeast and elucidated their functional dependencies with axial proteins Red1 and Hop1. At local levels, our analysis suggests that their axis-associated assembly involves two distinct pathways, Hop1-RMM-MRX and Red1-Spo11 core, whereas Spo11-core alone can bind DSB sites. Red1 also binds DSB sites in a DSB-dependent manner, supporting its critical role in DSB repair. At large scale, all ten DSB proteins and both axial proteins are enriched at DSB-hot domains 20-40 kb wide that correspond to sizes of the axis-loop units. Further analysis at short and long distances suggests that their local assembly, short-range/intra-loop distributions, and long-range/inter-loop distributions are distinctly controlled. Similarly in mouse spermatocytes, MEl4 and a Hop1 homolog HORMAD1 are enriched at DSB-hot domains 1-3 Mb wide, and their long-range distributions are distinctly controlled from their local assembly. Our results elucidate multi-layered mechanisms of chromosomal assembly of the DSB machinery in shaping the DSB landscape in yeast and mouse.

## Introduction

Three-dimensional chromosome structures control critical events in the nucleus such as transcription, DNA replication, and DNA repair. During meiosis, cells lose topologically associating domains (TADs) and organize specialized higher-order chromosome structure where loops of chromatin emanate from structural axes that are enriched with cohesin complexes, which hold sister chromatids together and extrude loops ^1–4^. Chromosome axes play critical roles in all processes of meiotic recombination including the formation and repair of programmed DNA double-strand breaks (DSBs). The repair of meiotic DSBs between homologous chromosomes (homologs) allows the reciprocal exchange of DNA contents between parental chromosomes to provide genetic diversity in the offspring. Each pair of homologs obtains at least one crossover, which ensures that chromosomes equally segregate into gametes ^5^.

In addition to cohesin, axial proteins that comprise axis-core proteins and HORMA (Hop1p, Rev7p, and MAD2)-domain-containing proteins organize chromosome axes. In budding yeast *Saccharomyces cerevisiae*, meiotic cohesin Rec8 and axial proteins Red1 and Hop1 are all required for full levels of DSBs and crossovers ^6–13^. Previous studies revealed the dependency relationships for their assembly at chromosome axes: Rec8 accumulates preferentially at the 3’ ends of convergent genes and recruits Red1 and Hop1 ^14–17^. While Red1 recruits Hop1 to Rec8 binding sites, Hop1 modulates Red1 distribution ^15, 18, 19^.

Red1 and Hop1 function as a basis of the assembly of proteins required for DSB formation (referred to as DSB proteins) on meiotic chromosomes. DSB proteins are conserved from yeasts to mammals, and ten proteins are required for DSB formation and form three distinct complexes in *S. cerevisiae* ^20^. Spo11 and topoisomerase VlB-like protein Rec102, which have structural similarities to the A subunit and the transducer domain of B subunit of the archaeal topoisomerase Vl (TopoVl), respectively ^21, 22^, form a Spo11 core complex with Ski8 and Rec104 ^23–30^. Mer2 forms an RMM complex with Rec114 and Mei4, all of which condense on DNA ^31^. Mer2 is phosphorylated by cyclin-dependent kinase (CDK) and Dbf4-dependent kinase (DDK), which is a prerequisite for the formation of the RMM complex on meiotic chromatin and for DSB formation ^17, 32–36^. The MRX complex comprising Mre11, Rad50, and Xrs2 (Nbs1 in other species) is essential for DSB formation specifically in *S. cerevisiae* ^37, 38^. Interactions between the three complexes allow the full assembly of the DSB proteins ^25, 31, 39–41^. Genome-wide chromatin immunoprecipitation (ChlP) analysis revealed that RMM components predominantly localized to chromosome axes in a Hop1/Red1-dependent manner ^17^. Importantly, fission yeast Rec7-Rec24-Rec15 (SFT) complex and mouse REC114-MEl4-lHO1 (RMl) complex (orthologs of Rec114-Mei4-Mer2) were also enriched at chromosome axes ^42–44^. Immunostaining foci of RMl on chromosome axes were largely abolished in *Hormad1^-/-^* mouse spermatocytes (HORMAD1 is a Hop1 homolog) ^42, 45^. Physical interactions of budding yeast Mer2, fission yeast Rec15, and mouse IHO1 with Hop1, Hop1, and HORMAD1, respectively ^42, 46, 47^, suggest the conserved role of Mer2-Hop1/Rec15-Hop1/lHO1-HORMAD1 interactions in recruiting the RMM/SFT/RMl complex to chromosome axes.

DSBs preferentially form at hotspots, which are located in nucleosome-depleted regions in chromatin loops ^48–50^ and tethered to chromosome axes ^6, 17^. DSB distribution is proposed to be controlled by multi-layered mechanisms from fine-scale chromatin structure including histone modifications to large-scale chromosome structure ^10, 50–55^. A prominent feature of large-scale DSB regulation is the generation of larger DSB-hot domains. Genome-wide analysis revealed that high ChlP density of Red1 and Hop1 on the three shortest chromosomes coincide with a high density of DSBs measured by sequencing of Spo11 oligos, byproducts of meiotic DSBs ^50^, and crossovers ^15, 17^. The overrepresentation of Red1 on the three shortest chromosomes was compromised in *hop1Δ* mutants despite Red1 being still enriched at Rec8-binding sites ^15^. Moreover, ChlP density of Rec114 and Mer2 was also higher on the three shortest chromosomes, and the overrepresentation of Rec114 was lost in both *hop1Δ* and *red1Δ* mutants ^10^. Accordingly, in *hop1Δ* or *red1Δ* mutants, genome-wide DSB levels were greatly reduced, and DSB density on the three shortest chromosomes decreased to the levels comparable to the other longer chromosomes ^10^. Thus, proper assembly of Red1, Hop1, Rec114, and Mer2 on chromosome axes makes the shortest chromosomes DSB-hot, which in turn ensures efficient crossovers and accurate segregation of homologs at meiosis I.

In contrast to whole chromosome-scale DSB regulation ^10, 15, 17^, our knowledge about DSB regulation at chromosomal domain scale is limited to the presence of DSB-hot domains 30-50 kb wide where DSB and Red1 densities are higher on the shortest chromosome III ^6^ and a weak anti-correlation between DSB and Rec8 densities on a few chromosomes ^50^. At local levels, beside extensive analysis of RMM components ^10, 17^, the full view of chromosomal assembly of the DSB machinery remains unsolved.

Here we provide a comprehensive analysis of the spatial distributions of DSB and axial proteins in yeast and elucidate their functional dependencies. Our results suggest that local assembly of the DSB machinery at chromosome axes involves two distinct pathways, Hop1-RMM-MRX and Red1-Spo11 core. Scale-dependent correlation between ChlP density of DSB and axial proteins and DSB density revealed that all DSB and axial proteins are enriched at DSB-hot large chromosomal domains that correspond to sizes of the axis-loop units, and suggests distinct mechanisms that regulate their distributions at short versus long distances. Dependency analysis supports distinct regulation between short and long distances, and further revealed that their chromosomal domain-scale distributions require protein interactions that are not involved in their local assembly. Moreover, we provide evidence that conserved DSB and axial proteins in mice are also enriched at DSB-hot large chromosomal domains and their chromosomal domain-scale distributions are distinctly regulated from their local assembly. Our findings thus demonstrate multi-layered mechanisms of chromosomal assembly of the DSB machinery in the context of tethered loop-axis complex in controlling DSB distribution in yeast and mice.

## Results

### All DSB proteins in budding yeast localize to chromosome axes with earlier association of Spo11-core components to DSB hotspots

To comprehensively understand the spatial distributions of the DSB proteins in budding yeast, we performed ChlP-seq for C-terminally FLAG- or HA-tagged DSB proteins at 3 hr and 4 hr after meiotic induction, when DSBs start to form and peak, respectively. At 3 hr, all Spo11 core and MRX components localized to chromosome axis sites, previously defined by ChlP-seq for Rec8-FLAG ^16^, like RMM components ^10, 17^ (**Fig. 1a, b, and Supplementary Fig. 1a, b**). Enrichment of DSB proteins at chromosome axis sites showed a high correlation, with Pearson’s *r* = 0.59-0.95 (**Supplementary Fig. 2**). In addition, all four Spo11 core components were enriched at DSB hotspots, previously defined by Spo11-oligo mapping ^50^, with clear enrichment at those with higher Spo11 enrichment at 3 hr (**Fig. 1a, c**). In contrast, MRX components were enriched at those DSB hotspots at 4 hr, but not 3 hr, like Rec114 (**Fig. 1a, c, and Supplementary Fig. 1a, c**). Enrichment of Spo11 core components at 3 hr and Spo11, Ski8, Rec114, Mre11, and Rad50 at 4 hr at DSB hotspots showed a high correlation, with Pearson’s *r* = 0.60-0.94 and 0.61-0.89, respectively (**Supplementary Fig. 3a, b**). These data indicate that all Spo11 and MRX components localize to chromosome axes like RMM components, and Spo11 core components localize to DSB hotspots earlier than RMM and MRX components.

**Fig. 1.**
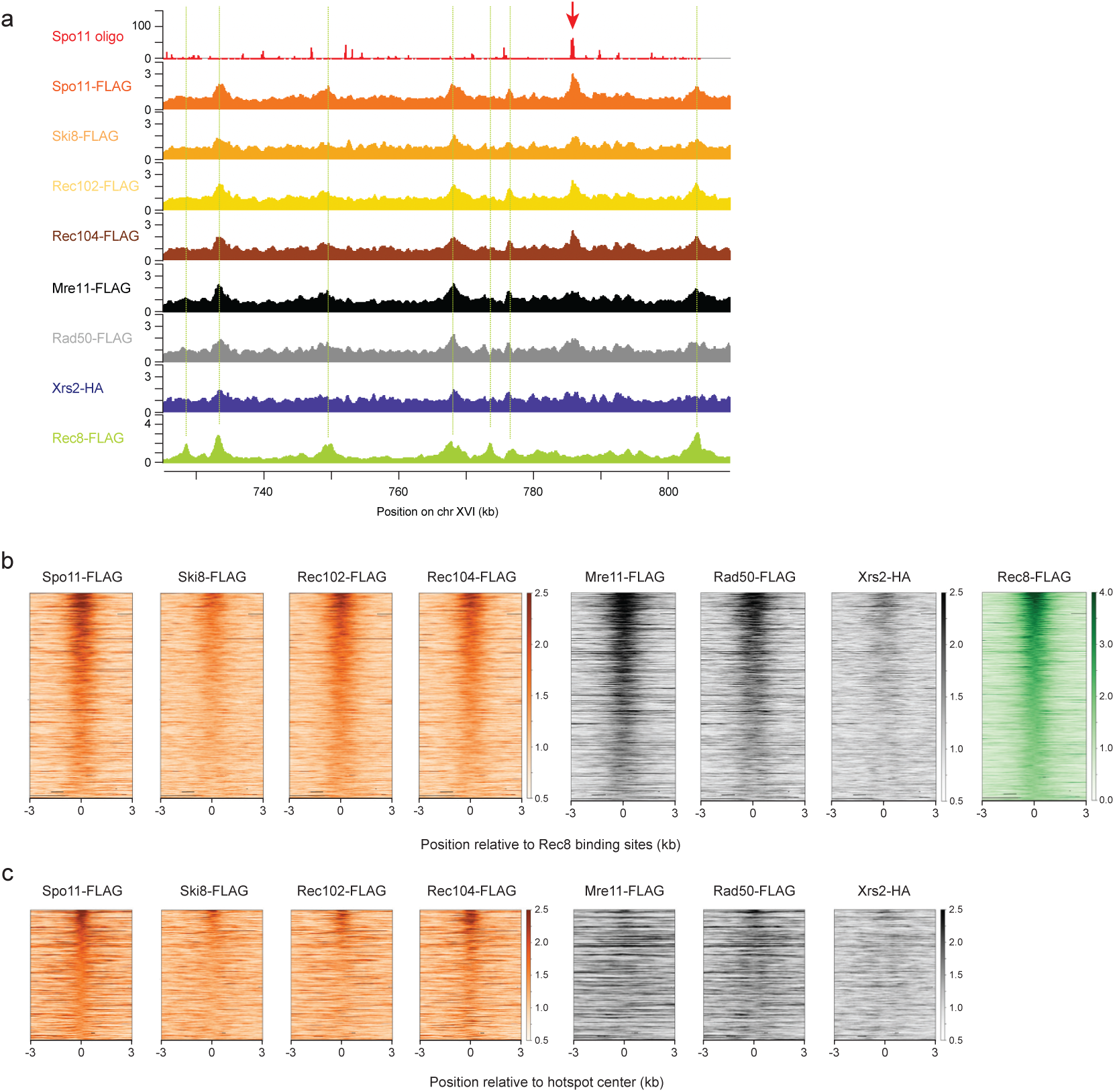
Spo11-core and MRX components localize to chromosome axes with additional binding of Spo11-core components at DSB hotspots. (a) ChIP signals of Spo11-core and MRX components in wild type at 3 hr after meiotic induction at a representative region on chromosome XVI. Spo11-oligo sequencing data from a published study ^76^ and ChIP signals of Spo11-FLAG and Rec8-FLAG in wild type at 3 hr and 4 hr after meiotic induction, respectively, from our previous study ^16^ are also shown. A red arrow and green dashed lines indicate a DSB hotspot with prominent Spo11 enrichment and Rec8 binding sites, respectively. (b) Heatmaps of ChIP signals of Spo11-core and MRX components in wild type at 3 hr after meiotic induction around chromosome axes. The top 540 sites with highest Rec8 enrichment in wild type at 3 hr among 724 Rec8 binding sites identified in our previous study ^16^ are ordered by Rec8 enrichment. ChIP signals are centered relative to Rec8 binding sites. (c) Heatmaps of ChIP signals of Spo11-core and MRX components in wild type at 3 hr after meiotic induction around DSB hotspots. The top 360 sites with highest Spo11 enrichment in wild type at 3 hr among 3,600 DSB hotspots identified in a previous study ^50^ are ordered by Spo11 enrichment. ChIP signals are centered relative to hotspot centers.

### Mre11 localization to chromosome axes depends on DDK-dependent Mer2 phosphorylation

While molecular mechanisms of Mre11 binding to DSB hotspots have been studied previously ^39, 41^, how axis localization of the MRX complex is regulated remain unsolved. Mer2 undergoes CDK-dependent phosphorylation followed by DDK-dependent phosphorylation, and CDK-dependent phosphorylation of Mer2 is critical for its interaction with Xrs2 by yeast two-hybrid assay ^34–36^. To explore the role of DDK-dependent Mer2 phosphorylation in axis localization of MRX components, we analyzed *mer2^S^*^11,15,19,22^,*^29A^* mutants (here after referred to as *mer2*^5x^*^SA^*) where five serine residues phosphorylated by DDK were substituted by alanine ^35^. While *mer2*^5x^*^SA^* mutation had little effect on Mer2 localization to chromosome axes, it reduced axis localization of Mei4 (**Fig. 2a-c**). Similarly, Mre11 binding to chromosome axes was greatly reduced in *mer2*^5x^*^SA^* cells. Importantly, axis localization of Mre11 was largely maintained in *spo11Δ* mutants (**Fig. 2a-c**), where Mre11 binding to DSB hotspots was reduced ^39^, indicating that altered Mre11 localization in *mer2*^5x^*^SA^* cells is not an indirect consequence of loss of DSBs. We conclude that axis localization of Mre11, and likely the full MRX complex, is dependent on DDK-dependent Mer2 phosphorylation and is distinctly regulated from its localization to DSB hotspots.

**Fig. 2.**
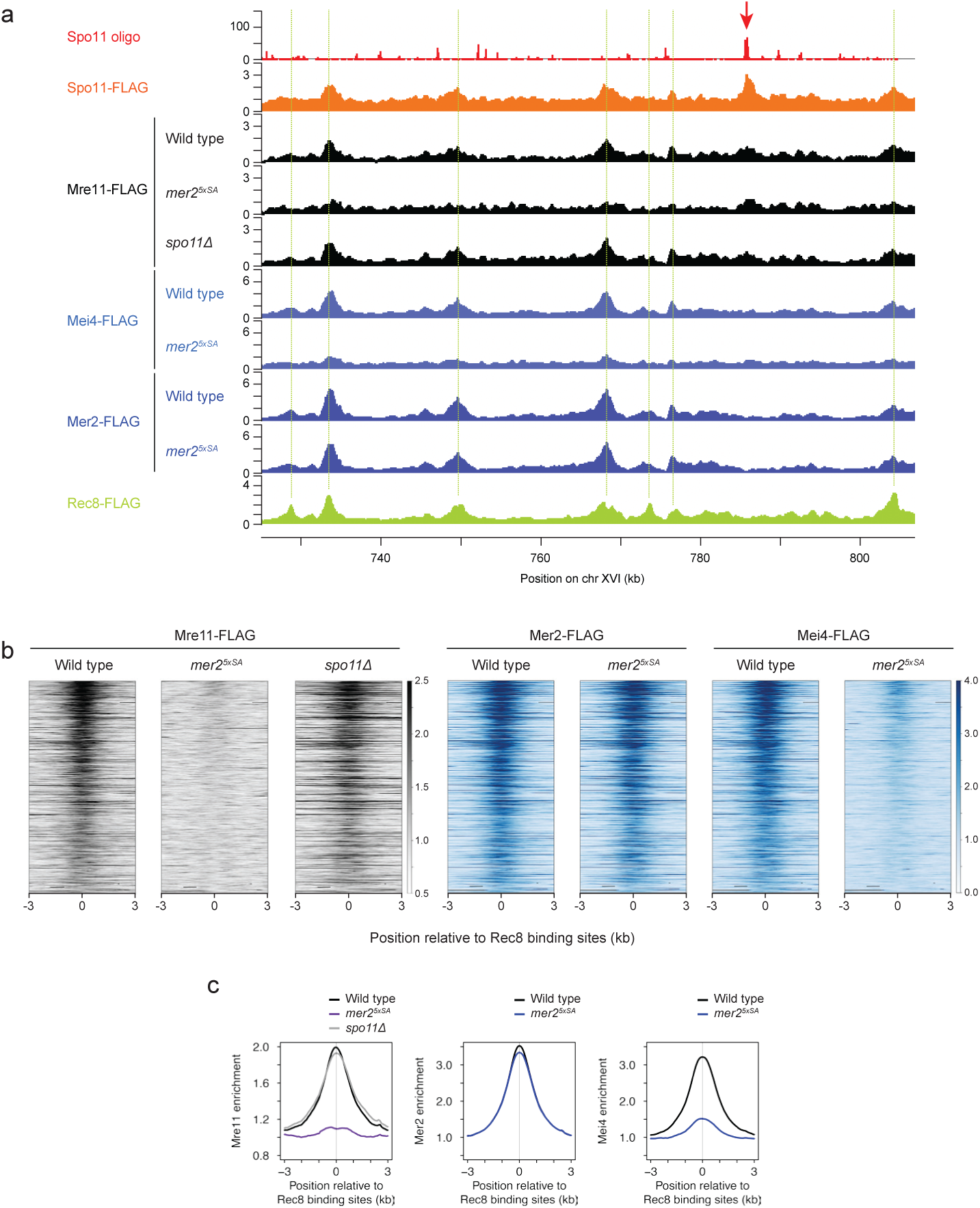
DDK-dependent Mer2 phosphorylation regulates axis assembly of MRX and RMM components. (a) ChIP signals of Mre11-FLAG, Mei4-FLAG, and Mer2-FLAG in wild type and indicated mutants at 3 hr after meiotic induction at the same chromosomal region shown in Fig. 1a. Spo11-oligo sequencing data and ChIP signals of Spo11-FLAG and Rec8-FLAG in wild type are reproduced from Fig. 1a to aid comparison. A red arrow and green dashed lines indicate a DSB hotspot with prominent Spo11 enrichment and Rec8 binding sites, respectively. (b) Heatmaps of ChIP signals of Mre11-FLAG, Mei4-FLAG, and Mer2-FLAG in wild type and indicated mutants at 3 hr after meiotic induction around chromosome axes, displayed as in Fig. 1b. (c) Metaplots of ChIP signals of Mre11-FLAG, Mer2-FLAG, and Mei4-FLAG around chromosome axis sites. Smoothed ChIP signals in wild type and indicated mutants at 3 hr after meiotic induction around ±3 kb of the 540 Rec8 binding sites used for heatmap representation in (b) are shown.

### Spo11 localization to chromosome axes is largely independent of RMM components and Hop1 but dependent on Red1

To understand the mechanisms that regulate axis localization of the Spo11 core complex, we analyzed Spo11 distribution in several mutants. In *mer2Δ* and *rec114Δ* mutants, Spo11 binding to chromosome axes was reduced but largely retained, in sharp contrast to the *rec102Δ* mutant, where axis localization of Spo11 was severely impaired (**Fig. 3a-c**). In *hop1Δ* mutants, where axis localization of RMM components was diminished ^17^, Spo11 binding to chromosome axes was reduced but retained at a similar level to *mer2Δ* and *rec114Δ* mutants (**Fig. 3a-c**). In the *red1Δ* mutant, in contrast, Spo11 binding to chromosome axes was nearly abolished (**Fig. 3a-c**). These results indicate that axis binding of Spo11, likely representing the Spo11 core complex, occurs in a Red1-dependent manner, but does not require Hop1 and the RMM complex; axis assembly of Hop1 and RMM components seem to rather facilitate efficient binding of the Spo11 core complex at chromosome axes.

**Fig. 3.**
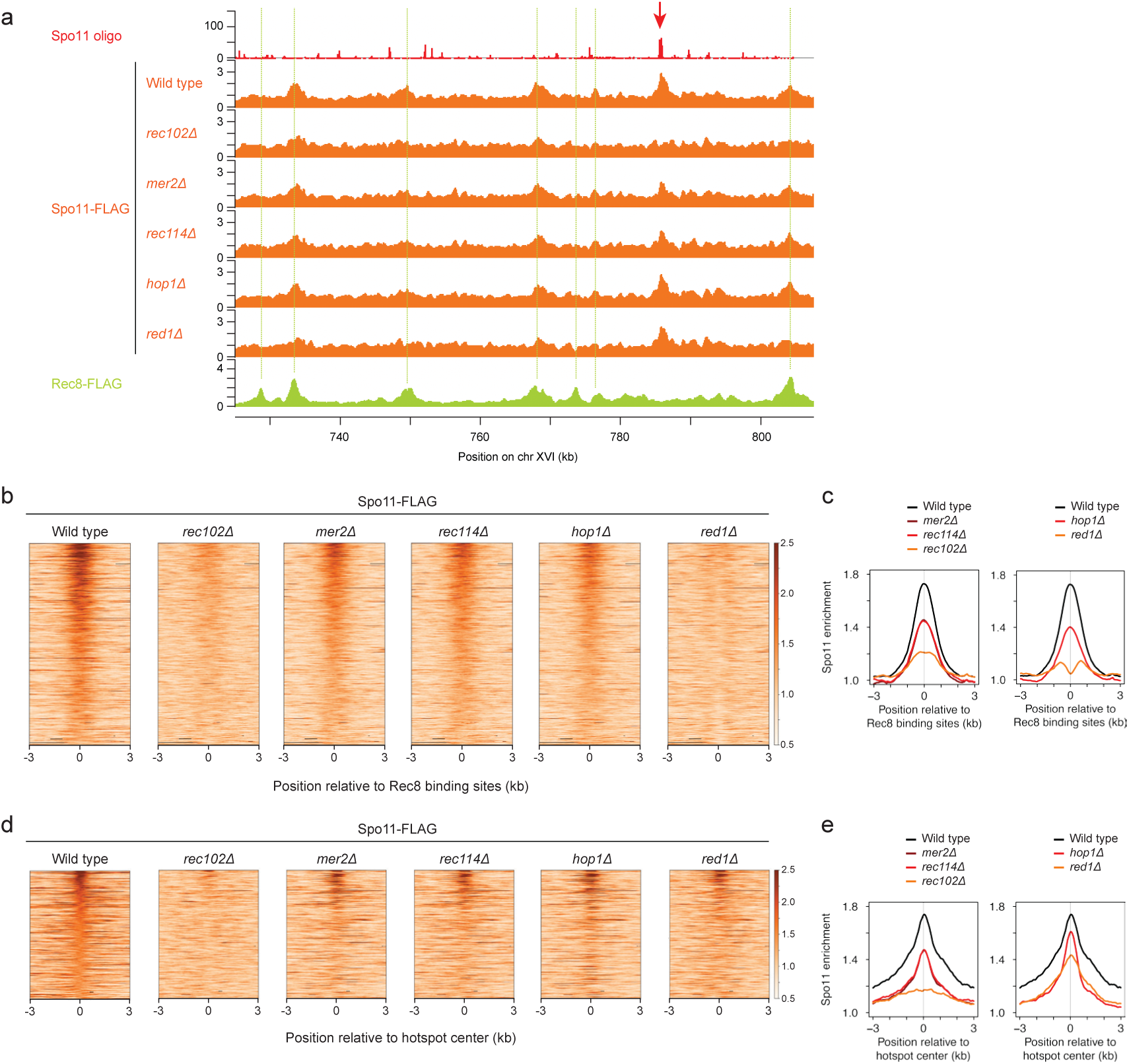
Spo11 localization to chromosome axes and DSB hotspots are distinctly regulated. (a) ChIP signals of Spo11-FLAG in wild type and indicated mutants at 3 hr after meiotic induction at the same chromosomal region shown in Fig. 1a. Spo11-oligo sequencing data and ChIP signals of Spo11-FLAG and Rec8-FLAG in wild type are reproduced from Fig. 1a to aid comparison. A red arrow and green dashed lines indicate a DSB hotspot with prominent Spo11 enrichment and Rec8 binding sites, respectively. (b) Heatmaps of ChIP signals of Spo11-FLAG in wild type and indicated mutants at 3 hr after meiotic induction around chromosome axes, displayed as in Fig. 1b. (c) Metaplots of ChIP signals of Spo11-FLAG around chromosome axis sites. Smoothed ChIP signals in wild type and indicated mutants at 3 hr after meiotic induction around ±3 kb of the 540 Rec8 binding sites used for heatmap representation in (b) are shown. (d) Heatmaps of ChIP signals of Spo11-FLAG in wild type and indicated mutants at 3 hr after meiotic induction around DSB hotspots, displayed as in Fig. 1c. (e) Metaplots of ChIP signals of Spo11-FLAG around DSB hotspots. Smoothed ChIP signals in wild type and indicated mutants at 3 hr after meiotic induction around ±3 kb of the 360 DSB hotspots used for heatmap representation in (d) are shown.

### Spo11 localization to DSB hotspots is largely independent of RMM components and axial proteins but dependent on Rec102

In *mer2Δ*, *rec114Δ*, *hop1Δ*, and *red1Δ* mutants, Spo11 binding to DSB hotspots was reduced but retained (**Fig. 3a, d, e**). In contrast, Spo11 binding to DSB hotspots was nearly lost in the *rec102Δ* mutant (**Fig. 3a, d, e**). This indicates that, like axis localization, Spo11 binding to DSB hotspots occurs in the absence of RMM components and Hop1, but requires Rec102. Notably, loss of Spo11 binding from chromosome axes, but not DSB hotspots in the *red1Δ* mutant indicates that Spo11 binding to chromosome axes and DSB hotspots are distinctly regulated.

### Red1 binds crossover-designated sites in a DSB-dependent manner

While our previous ChlP-seq for C-terminally FLAG-tagged Red1 at 3 hr showed its predominant localization to chromosome axes ^56^ consistent with other published data ^15, 17^, weak enrichment of Red1-FLAG at some DSB hotspots led us to analyze Red1 distribution at later timepoints. Red1-FLAG enrichment at DSB hotspots, which is weakly observed from 3 hr, was evident at 4 hr and further enhanced at 5 hr (**Fig. 4a-c**). In contrast, Red1-FLAG enrichment at chromosome axes peaked at 3 hr, and decreased as meiotic prophase l progressed (**Fig. 4a, d, e**). Notably, Red1-FLAG enrichment at some DSB hotspots was higher than that of adjacent chromosome axis sites at later timepoints (**Fig. 4a**), supporting its critical role in inter-homolog DSB repair ^7, 57^.

**Fig. 4.**
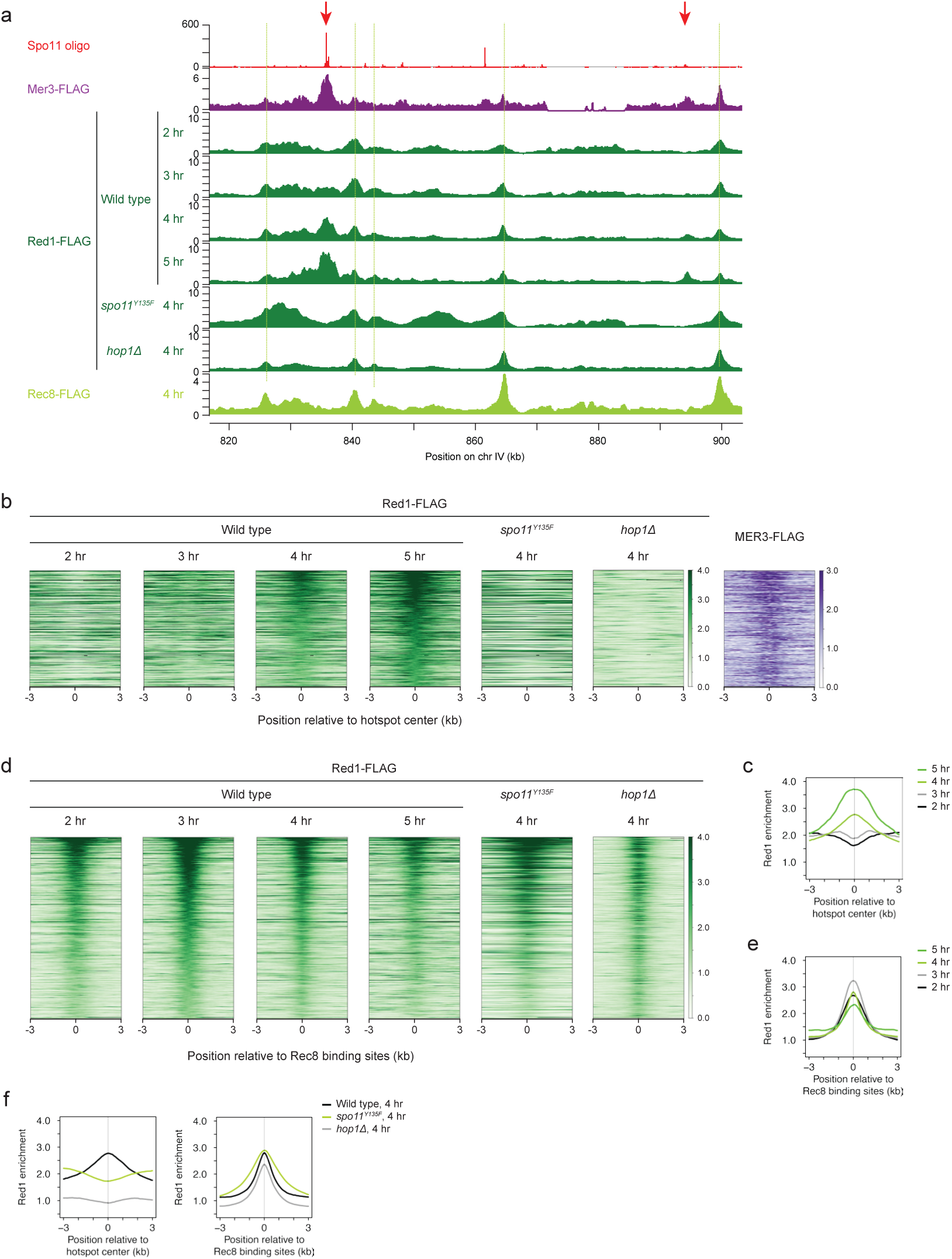
Red1 localizes to crossover-designated sites in a DSB- and Hop1-dependent manner in addition to chromosome axes. (a) ChIP signals of Red1-FLAG in wild type at indicated timepoints after meiotic induction at a representative region on chromosome IV. A different region from Fig. 1-3 is shown as DSB hotspots within the chromosomal region represented in earlier figures show no clear Red1-FLAG enrichment. Spo11-oligo sequencing data and ChIP signals of Rec8-FLAG and Mer3-FLAG in wild type at 4 hr after meiotic induction from previous studies ^16, 58, 76^ are also shown. Red arrows and green dashed lines indicate DSB hotspots with prominent Red1 enrichment at later timepoints and Rec8 binding sites, respectively. (b) Heatmaps of ChIP signals of Red1-FLAG and Mer3-FLAG in wild type and indicated mutants at indicated timepoints after meiotic induction around DSB hotspots. The top 360 sites with highest Red1 enrichment in wild type at 5 hr among 3,600 DSB hotspots identified in a previous study ^50^ are ordered by Red1 enrichment. ChIP signals are centered relative to hotspot centers. (c) Metaplots of ChIP signals of Red1-FLAG around DSB hotspots. Smoothed ChIP signals in wild type at indicated time points after meiotic induction around ±3 kb of the 360 DSB hotspots used for heatmap representation in (b) are shown. (d) Heatmaps of ChIP signals of Red1-FLAG in wild type and indicated mutants at indicated timepoints after meiotic induction around chromosome axes. The top 540 sites with highest Red1 enrichment in wild type at 3 hr among 724 Rec8 binding sites identified in our previous study ^16^ are ordered by Red1 enrichment. ChIP signals are centered relative to Rec8 binding sites. (e) Metaplots of ChIP signals of Red1-FLAG around chromosome axis sites. Smoothed ChIP signals in wild type at indicated time points after meiotic induction around ±3 kb of the 540 Rec8 binding sites used for heatmap representation in (d) are shown. (f) Metaplots of ChIP signals of Red1-FLAG around DSB hotspots (left) and chromosome axis sites (right). Smoothed ChIP signals in wild type and indicated mutants at 4 hr after meiotic induction around ±3 kb of the 360 DSB hotspots and 540 Rec8 binding sites used for heatmap representation in (b) and (d), respectively, are shown.

Red1-FLAG enrichment at DSB hotspots showed a high correlation with Mer3, a pro-crossover factor that preferentially binds crossover-designated sites ^58^ , but only a weak correlation with Spo11 (**Supplementary Fig. 4a, b**; Pearson’s *r* = 0.62 and 0.61 with Mer3 and 0.27 and 0.24 with Spo11 at 4 hr and 5 hr, respectively). This is in sharp contrast to DSB proteins, which showed a high correlation with Spo11 at 4 hr (**Supplementary Fig. 3b**). In the *spo11^Y^*^135^*^F^* mutant, Red1-FLAG binding to DSB hotspots was diminished while its binding to chromosome axes was enhanced (**Fig. 4a, b, d, f**). Thus, axial Red1 preferentially binds crossover-designated sites in a DSB-dependent manner.

### Dynamic changes in Red1 distribution during early prophase I is controlled by DSB formation and Hop1

Altered Red1 distribution in the *spo11^Y^*^135^*^F^* mutant was also seen in other chromosomal regions than DSB hotspots and chromosome axes: Red1-FLAG was enhanced outside chromosome axes, showing broad peaks spanning >10 kb wide at some regions (**Supplementary Fig. 5a**). These regions included, but were not restricted to, previously identified Rec8-independent Red1 binding domains, referred to as islands ^15, 18^ (**Supplementary Fig. 5a, b**). Other DSB-defective mutants *spo11Δ*, *rec104Δ*, *mer2*^5x^*^SA^*, *mre11Δ*, and *rad50Δ* also showed altered Red1 distribution like the *spo11^Y^*^135^*^F^*mutant (**Supplementary Fig. 5a-d, 6**). In contrast, Red1-FLAG binding to DSB hotspots was abolished but its binding to chromosome axes and islands was not increased in the *hop1Δ* mutant (**Fig. 4a, b, d, f, and Supplementary Fig. 5a, b**). These data indicate the dynamic nature of Red1 distribution on early prophase-l chromosomes, which is controlled by DSB formation and Hop1.

### DSB and axial proteins are enriched at DSB-hot chromosomal domains with distinct correlations with DSB density at short and long distances

As described in the Introduction, large-scale chromosome structure plays critical roles in controlling DSB distribution ^6, 10, 15, 17, 56^ . To gain mechanistic insights into DSB regulation at chromosomal domain scale, we examined correlations between ChlP density of DSB and axis-organizing proteins and Spo11-oligo density (i.e. DSB density) at variable window sizes ranging 0.5-40 kb in a way described in previous studies ^50, 56^ with a comprehensive data set of their spatial distributions.

ChlP density of Red1 at 3 hr showed a scale-dependent correlation with DSB density (**Fig. 5a**). Red1 density was weakly anti-correlated with DSB density for small windows (≤ 1 kb), but had a positive correlation with DSB density at larger windows. Correlations between Red1 and DSB densities showed a sharp, nearly linear increase up to 7.5 kb, followed by a gradual, curve increase for window sizes > 7.5 kb, which reached nearly a plateau for window sizes > 20-kb with Pearson’s *r* = 0.76 for data assessed in a 40-kb window (**Fig. 5a**). ChlP density of Hop1 in a published study ^15^ showed a similar pattern in correlations with DSB density to Red1, with Pearson’s *r* = 0.69 at a 40-kb window (**Fig. 5a**). This supports a previous finding that DSB-hot chromosomal domains 30-50 kb wide are enriched for Red1 on chromosome III ^6^ and further suggests distinct mechanisms of DSB regulation at short (≤ 7.5 kb) and long (> 7.5 kb) distances. Rec8 density was anti-correlated with DSB density for window sizes ≤ 5 kb with a small peak at a 1-kb window and showed a positive correlation with DSB density for window sizes > 5 kb (**Fig. 5a**). This correlation was lower than Red1 and Hop1 over all window sizes analyzed with Pearson’s *r* = 0.50 at a 40-kb window, consistent with distinct distributions of Rec8 from axial proteins at chromosomal domain scale ^6, 56^.

**Fig. 5.**
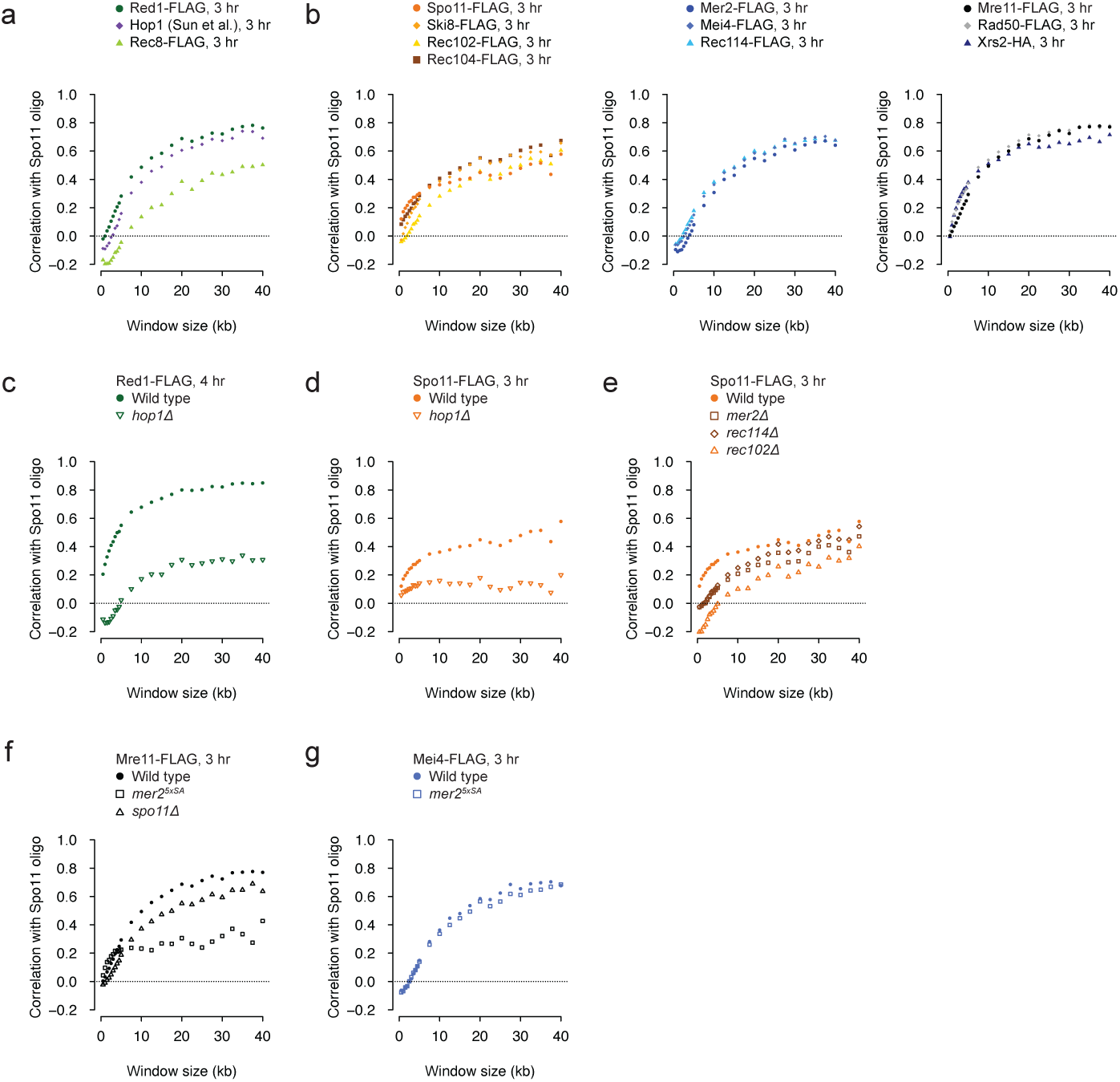
ChIP density of DSB and axis-organizing proteins shows scale-dependent correlations with DSB frequency at chromosomal domain scale in budding yeast. (a) Scale-dependent correlations of DSB frequency with ChIP density of axial proteins and Rec8-FLAG at chromosomal domain scale. Spo11-oligo density in wild type from a published study ^76^ and ChIP signals of indicated proteins in wild type at 3 hr after meiotic induction in non-overlapping bins of varying sizes were compared and Pearson’s *r* values were plotted. 20 kb of each chromosomal end, centromeric region spanning 20 kb of each chromosome, and rDNA cluster chromosome XII were excluded. Hop1 ChIP data is from a published study ^15^. (b) Scale-dependent correlations of DSB frequency with ChIP density of indicated DSB proteins at 3 hr after meiotic induction, displayed as in (a) (c-g) Scale-dependent correlations of DSB frequency with ChIP density of Red1-FLAG (c), Spo11-FLAG (d, e), Mre11-FLAG (f), and Mei4-FLAG (g) in wild type and indicated mutants at indicated time points after meiotic induction, displayed as in (a).

All ten DSB proteins showed scale-dependent correlations with DSB density like axial proteins. A sharper increase in these correlations for short windows (∼7.5 kb) was followed by a more gradual increase for large windows, reaching nearly a plateau for window sizes > 20-kb with Pearson’s *r* = 0.58-0.67, 0.64-0.68, and 0.71-0.78 for Spo11-core, RMM, and MRX components, respectively, at 3 hr at 40-kb windows (**Fig. 5b**). This indicates that DSB-hot chromosomal domains are enriched for not only axial proteins but also all DSB proteins. For short windows (∼7.5 kb), most of Spo11-core and MRX components showed a higher positive correlation with DSB density than RMM components (**Fig. 5b**). In contrast, with large windows, RMM components showed a high positive correlation with DSB density that were comparable to Spo11-core and MRX components (**Fig. 5b**). Similar patterns were also observed at 4 hr (**Supplementary Fig. 7a**). This supports the idea that mechanisms of DSB regulation are distinct between short (≤ 7.5 kb) and long (> 7.5 kb) distances: the amount of the Spo11-core and MRX complexes may be more important at short chromosomal domains whereas that of all three complexes seem critical at long chromosomal domains.

### Coordinated enrichment of Red1, Spo11 and DSBs at large chromosomal domains requires Hop1

To understand the underlying mechanisms that regulate distributions of the DSB and axial proteins at chromosomal domain scale, we first analyzed *hop1Δ* mutants, where DSB formation is severely impaired and DSB distribution is altered ^10^. When compared to DSB density in wild type, Red1 density in the *hop1Δ* mutant at 4 hr showed a weak anti-correlation and positive correlation for window sizes ≤ 4.5 kb and > 4.5 kb, respectively, with Pearson’s *r* = 0.31 at a 40-kb window, in sharp contrast to a high positive correlation with Pearson’s *r* > 0.7 for window sizes > 10 kb in wild type (**Fig. 5c**). This correlation was even lower than other DSB-defective mutants, all of which showed stinkingly similar patterns, over all window sizes analyzed (**Supplementary Fig. 7b**; Pearson’s *r* =0.58, 0.59, 0.62, 0.58, 0.67, and 0.65 in *spo11^Y^*^135^*^F^*, *spo11Δ*, *rec104Δ*, *mer2*^5x^*^SA^*, *mre11Δ*, and *rad50Δ*, respectively, at 4 hr at 40-kb windows). Spo11 density in the *hop1Δ* mutant at 3 hr showed a weak positive correlation with Pearson’s *r* < 0.2, which was lower than wild type over all window sizes analyzed, like Red1 (**Fig. 5d**). This indicates that high enrichment of Red1 and Spo11 at DSB-hot chromosomal domains requires Hop1.

When compared to DSB density in the *hop1Δ* mutant ^10^, Red1 density in wild type showed a largely scale-independent weak positive correlation with Pearson’s *r* < 0.2 while that in the *hop1Δ* mutant showed a scale-dependent correlation with Pearson’s *r* = 0.47 at a 40-kb window (**Supplementary Fig. 7c**). Spo11 density in both wild type and the *hop1Δ* mutant showed a largely scale-independent weak positive correlation with Pearson’s *r* < 0.4 (**Supplementary Fig. 7d**). Thus, chromosomal domains enriched for Red1 and Spo11 in wild type are not DSB-hot in the absence of Hop1. These results indicate that Hop1 is required for coordinated enrichment of Red1, Spo11, and DSBs at large chromosomal domains: Hop1 generates DSB-hot chromosomal domains by controlling Red1 and Spo11 distributions.

Notably, Spo11 density in the *red1Δ* mutant at 3 hr showed nearly wild-type levels of positive correlation with wild-type DSB density, despite a stronger reduction of its axis localization in *red1Δ* than *hop1Δ* mutants (**Supplementary Fig. 7e**; Pearson’s *r* = 0.58 and 0.48 in wild type and *red1Δ,* respectively, at 3 hr at 40-kb windows). This suggests that Spo11 distribution at chromosomal domain scale is distinctly regulated from its local axis assembly.

### Distributions of Spo11 and Mre11 at long distance are distinctly controlled from their distributions at short distance and local axis assembly

We next addressed the impacts of other DSB-defective mutations described above on the spatial distributions of DSB proteins at chromosomal domain scale. Given no DSB formation in these mutants, we compared their distributions in mutants with wild-type DSB density at variable window sizes ranging 0.5-40 kb to assess changes in their distributions.

Spo11 density in *mer2Δ*, *rec114Δ*, and *rec102Δ* mutants showed correlations that were reduced over all window sizes analyzed compared to wild type, to the greatest degree in the *rec102Δ* mutant (**Fig. 5e**). While levels of the reduction were largely scale-independent in the *rec102Δ*, a greater reduction was observed for window sizes ≤ 10 kb in *mer2Δ* and *rec114Δ* mutants (**Fig. 5e**; Pearson’s *r* = 0.36, 0.21, 0.25, and 0.10 in wild type, *mer2Δ, rec114Δ*, and *rec102Δ*, respectively, at 3 hr at 10-kb windows).

Mre11 also showed distinct patterns in changes in correlations with DSB density in mutants. While correlations between Mre11 and DSB densities were nearly comparable to wild type over all window sizes analyzed in the *spo11Δ* mutant, they were prominently reduced for window sizes ≥ 10 kb in the *mer2*^5x^*^SA^* mutant (**Fig. 5f and Supplementary Fig. 7f**; Pearson’s *r* = 0.77, 0.64, and 0.43 in wild type, *spo11Δ*, *mer2*^5x^*^SA^*, respectively, at 3 hr at 40-kb windows). Notably, the *mer2*^5x^*^SA^* mutation had little impacts on correlations of not only Mer2 but also Mei4 with DSB density, despite a strong reduction of axis localization of Mei4 like Mre11 in *mer2*^5x^*^SA^* cells (**Fig. 5g and Supplementary Fig. 7g**). This clearly indicates that mechanisms that facilitate local axis assembly and that regulate chromosomal domain-scale distribution are distinct. Furthermore, distinct patterns in changes in Spo11 and Mre11 distributions at short versus long distances in mutants suggest that their distributions are distinctly regulated at short and long chromosomal domains.

### DSB-hot chromosomal domains are enriched for HORMAD1, MEI4, and CTCF, but not SYCP3, IHO1, and cohesin in mouse spermatocytes

DSB and axis-organizing proteins are evolutionary conserved from yeasts to mammals ^20^. To gain insights into DSB regulation at chromosomal domain scale in mammals, we analyzed published ChlP-seq data in mouse spermatocytes ^59, 60^ and examined correlations between ChlP density of DSB and axis-organizing proteins and SPO11-oligo density (i.e. DSB density). Given that loop lengths in mouse autosomes are 50-fold larger than those in budding yeast ^16^ , we examined correlations at variable window sizes ranging 1 kb-3 Mb and displayed the results with two scales: ≤ 100 kb and ≤ 3 Mb (**Fig. 6a-h**).

**Fig. 6.**
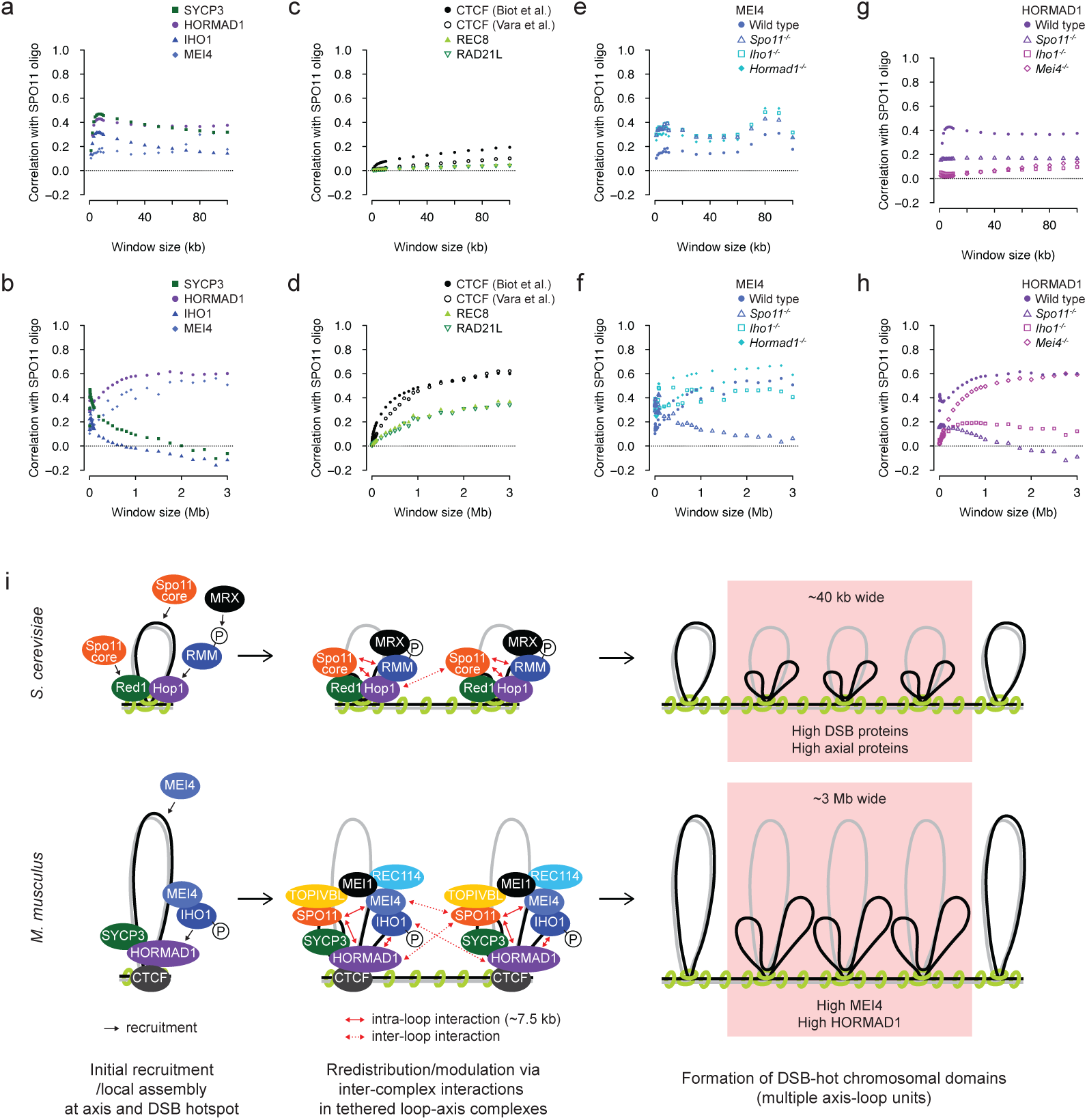
ChIP density of DSB and axis-organizing proteins shows scale-dependent correlations with DSB frequency at chromosomal domain scale in mouse spermatocytes. (a, b) Scale-dependent correlations of DSB frequency with ChIP density of DSB and axial proteins at short (a) and large (b) chromosomal domain scale. SPO11-oligo density and ChIP signals of indicated proteins in wild-type mouse spermatocytes from published studies ^59, 60, 75^ in non-overlapping bins of varying sizes up to 100 kb (a) and 3 Mb (b) were compared and Pearson’s *r* values were plotted. Sex chromosomes and 3 Mb from left end of each autosome were excluded. (c, d) Scale-dependent correlations of DSB frequency with ChIP density of CTCF and cohesin at short (c) and large (d) chromosomal domain scale, displayed as in (a) and (b), respectively. (e, f) Correlations of DSB frequency with ChIP density of MEI4 at short (e) and large (f) chromosomal domain scale in wild type and indicated mutants, displayed as in (a) and (b), respectively. (g-h) Correlations of DSB frequency with ChIP density of HORMAD1 at short (e) and large (f) chromosomal domain scale in wild type and indicated mutants, displayed as in (a) and (b), respectively. (i) A model of multi-layered control of chromosomal assembly of the DSB machinery and DSB distribution in the context of tethered loop-axis complexes in budding yeast and mice. On budding yeast chromosomes, axial proteins Red1 and Hop1 bind cohesin Rec8 binding sites and recruit DSB proteins in the three complexes to chromosome axes via two distinct pathways; Hop1-RMM-MRX and Red1-Spo11 core (black arrows). The Spo11 core complex alone can bind DSB hotspots in loops. The full assembly of the DSB machinery in the tethered loop-axis complexes further modulates their distribution via inter-complex interactions within (red arrows, ∼7.5 kb) and between (red dashed arrows) the axis-loop units, generating chromosomal domains ∼40 kb wide ranging multiple axis-loop units where all DSB and axial proteins are enriched and DSBs form with higher frequency. On mouse autosomes, axial proteins SYCP3 and HORMAD1 and DSB proteins MEl4 and IHO1 bind chromosome axes organized by cohesin REC8, RAD21L, and CTCF. HORMAD1 recruits IHO1 to chromosome axes, which requires DDK-dependent phosphorylation of IHO1 ^42, 66^. MEl4 likely can bind DSB hotspots in loops, and SYCP3, HORMAD1, and IHO1 bind DSB hotspots in the tethered loop-axis complexes ^59^. Whether SPO11, TOPVlBL, REC114, and MEl1 bind chromosome axes or DSB hotspots is currently unknown. ANKRD31, which is critical for DSBs on the pseudoautosomal region on sex chromosomes in mouse spermatocytes and regulates autosomal DSBs likely via distinct mechanisms from other DSB proteins ^45, 66, 77, 78^, is not shown. Initial recruitment and local assembly of each protein/complex via distinct pathways are followed by the full assembly of the DSB machinery in the tethered loop-axis complexes to modulates their distribution via inter-complex interactions. In mice, chromosomal domains ∼3 Mb wide with higher MEl4 and HORMAD1, but not SYCP3 and IHO1, form DSBs with higher frequency.

For window sizes ≤ 100 kb, SYCP3, HORMAD1, and IHO1 similarly showed a weak positive correlation with DSB density with a peak at 7-kb windows with Pearson’s *r* = 0.47, 0.43 and 0.32 for SYCP3, HORMAD1 and IHO1, respectively (**Fig. 6a**). This is consistent with nearly identical levels of their enrichment at resected DSB ends around DSB hotspots ^59^ (**Supplementary Fig. 8a**). However, with larger windows, patterns of their scale-dependent correlations with DSB density were distinct. HORMAD1 showed a positive correlation that was increased for window sizes ≥ 100 kb and remained high for window sized ≥ 1 Mb, with Pearson’s *r* = 0.58 and 0.60 at 1-Mb and 3-Mb windows, respectively (**Fig. 6b**). In contrast, SYCP3 and IHO1 showed a decrease in a positive correlation for window sizes ≥ 7 kb and eventually showed a weak negative correlation for windows > 2 and >1 Mb, with Pearson’s *r* = -0.062 and -0.12 at 3-Mb windows, respectively (**Fig. 6a, b**).

MEl4 showed a largely scale-independent positive correlation with DSB density for window sizes < 100 kb, which was followed by a gradual increase in positive correlations for window sizes ≥ 100 kb and remained high for window sizes > 2 Mb with Pearson’s *r* = 0.51 at a 3-Mb window (**Fig. 6a, b**). CTCF, REC8, and RAD21L showed an increase in a positive correlation that reached nearly a plateau for window sizes > 2 Mb (**Fig. 6c, d**). While a positive correlation of CTCF was comparable to HORMAD1 and MEl4 at large windows, that of REC8 and RAD21L was lower than those over all window sizes analyzed (**Fig. 6a-d**; Pearson’s *r* = 0.60, 0.62, 0.37, and 0.34 at 3-Mb windows for CTCF at leptonema/zygonema ^59^, CTCF at pachynema/diplonema ^60^, REC8, and RAD21L, respectively). These results indicate that DSB-hot chromosomal domains 2-3 Mb wide are enriched for HORMAD1, IHO1, and CTCF, but not for SYCP3, MEl4, REC8, and RAD21L.

### Local distributions of SYCP3, HORMAD1, and IHO1 around chromosome axes are distinct in mouse spermatocytes

A high positive correlation of HORMAD1, but not SYCP3 and IHO1, with DSB density at large chromosomal domains was unexpected, given their similar local enrichment at DSB hotspots, functional elements, and CTCF sites ^59^. To address whether they show distinct local distributions that were not described in a previous study ^59^, we reanalyzed published ChlP-seq data ^59, 60^. Given that CTCF functions with cohesin to organize chromosome structures in somatic cells ^61, 62^ and shows a linear immunostaining pattern on chromosome axes in pachytene spermatocytes in mice ^60^, we classified potential axis sites into three types: CTCF biding sites where neither REC8 or RAD21 is enriched (CTCF-only), CTCF binding sites where either REC8 or RAD21 is enriched (CTCF + cohesin), and cohesin binding sites where CTCF is little enriched (cohesin-only). Consistent with REC8 and RAD21L being enriched at gene regulatory elements including promoters and enhances ^60^, their binding sites overlapped with H3K4me3-enriched functional elements defined in a previous study ^59^ , while CTCF-only sites lacked H3K4me3 marks (**Supplementary Fig. 8a**).

SYCP3 and HORMAD1 showed similar distributions at cohesin-only sites and DSB hotspots. In contrast, our analysis revealed clear distinction of their distributions at CTCF-only and CTCF + cohesin sites. First, SYCP3 was enriched at around a half of CTCF-only sites and depleted from the other half, whereas HORMAD1 was enriched at most of CTCF-only sites, even at those where SYCP3 was depleted (**Supplementary Fig. 8a**). Second, while HORMAD1 was preferentially enriched at where cohesin REC8 and RAD21L strongly bind among CTCF-cohesin sites, SYCP3 enrichment at CTCF-cohesin sites showed little correlation with levels of cohesin binding (**Supplementary Fig. 8a, b**).

IHO1 showed similar distributions to SYCP3 and HORMAD1 at DSB hotspots and CTCF-only sites. However, IHO1 was depleted from most of CTCF-cohesin sites, preferentially from where cohesin REC8 and RAD21L strongly bind, in sharp contrast to HORMAD1 enrichment at those sites (**Supplementary Fig. 8a, b**). Among cohesin-only sites, IHO1 was also depleted from where cohesin strongly bind, while enriched at the rest (**Supplementary Fig. 8a, b**). Thus, despite a strong dependency of chromosomal localization of IHO1 on HORMAD1 and similar immunostaining patterns of SYCP3, HORMAD1, and IHO1 on unsynapsed chromosome axes ^42^, their distributions at chromosome axes are distinct.

### High enrichment of MEI4 and HORMAD1, but not CTCF, at DSB-hot chromosomal domains requires SPO11 in mouse spermatocytes

Our mutant analysis in budding yeast suggests that chromosomal domain-scale distributions of DSB and axial proteins are distinctly regulated from their local assembly. To address whether the similar mechanisms exist in mice, we analyzed published ChlP-seq data for MEl4 and HORMAD1 in mutants ^59^ . For window sizes ≤ 100 kb, changes in correlations between MEl4 and DSB densities reflected increased local enrichment of MEl4 at DSB hotspots ^59^ (**Supplementary Fig. 9a**): a positive correlation was similarly increased in *Iho1^-/-^*, *Spo11^-/-^*, and *Hormad1^-/-^*spermatocytes (**Fig. 6e**; Pearson’s *r* = 0.30, 0.48, 0.43, and 0.52 at 80-kb windows in wild type, *Iho1^-/-^*, *Spo11^-/-^*, and *Hormad1^-/-^*, respectively). In contrast, with large windows, MEl4 showed a weak positive correlation that was greatly lower than wild type only in *Spo11^-/-^* spermatocytes (**Fig. 6f**; Pearson’s *r* = 0.39, 0.37, 0.48, and 0,13 at 1-Mb windows and 0.51, 0.41, 0.59, and 0.062 at 3-Mb windows in wild type, *Iho1^-/-^*, *Hormad1^-/-^*, and *Spo11^-/-^*, respectively). This indicates that SPO11, but not DSBs, is critical for high enrichment of MEl4 at DSB-hot chromosomal domains.

HORMAD1 also showed distinct patterns in correlations with DSB density in mutants. For window sizes ≤ 100 kb, a positive correlation between HORMAD1 and DSB densities was lower than wild type in *Spo11^-/-^*, *Mei4^-/-^*, and *Iho1^-/-^*spermatocytes (**Fig. 6f**; Pearson’s *r* = 0.37, 0.17, 0.066 and 0,061 at 40-kb windows in wild type, *Spo11^-/-^*, *Mei4^-/-^*, and *Iho1^-/-^*, respectively). Levels of the reduction corresponded to those of local HORMAD1 enrichment at DSB hotspots, like MEl4 ^59^ (**Supplementary Fig. 9b**). With large windows, HORMAD1 in *Spo11^-/-^* and *Iho1^-/-^*, but not *Mei4^-/-^* spermatocytes, showed correlations that were reduced over all window sizes analyzed compared to wild type, to the greater degree in *Spo11^-/-^* spermatocytes (**Fig. 6h**; Pearson’s *r* = 0.58, 0.052, 0.19, and 0.49 at 1-Mb windows and 0.60, -0.091, 0.12 and 0.60 at 3-Mb windows in wild type, *Spo11^-/-^*, *Iho1^-/-^*, and *Mei4^-/-^*, respectively). Thus, not only SPO11 but also IHO1 is important for high HORMAD1 enrichment at DSB-hot chromosomal domains. Furthermore, reduced but retained local HORMAD1 enrichment at most of chromosome axis sites in *Iho1^-/-^* spermatocytes, in contrast to its depletion from cohesin binding sites in *Spo11^-/-^* spermatocytes (**Supplementary Fig. 9b**), suggests that chromosomal domain-scale distributions of DSB and axial proteins are also distinctly regulated from their local assembly in mice.

In contrast to MEl4 and HORMAD1, CTCF in *Spo11^-/-^* spermatocytes showed a positive correlation that was nearly comparable to wild type over all window sizes analyzed (**Supplementary Fig. 9c**; Pearson’s *r* = 0.60 and 0.51 in wild type and *Spo11^-/-^* at 3-Mb windows, respectively). Thus, SPO11 has little impact on CTCF distribution.

## Discussion

### Two distinct pathways, Hop1-RMM-MRX and Red1-Spo11 core, collaborate for the local assembly of the full DSB machinery at chromosome axes in budding yeast

Our comprehensive genome-wide ChlP-seq analysis in budding yeast revealed that all ten DSB proteins localized to chromosome axes. Further mutant analysis suggests that axis localization of MRX and Spo11 core complexes depends on DDK-dependent Mer2 phosphorylation and Red1, respectively. Given that axis localization of RMM components depends on Hop1 ^15, 17, 46^, we propose that two distinct pathways, Hop1-RMM-MRX and Red1-Spo11 core, collaborate to recruit the full DSB machinery at chromosome axes (**Fig. 6i**).

Earlier association of Spo11 core components at DSB hotspots than RMM and MRX components and Spo11 binding at DSB hotspots in the absence of RMM components and axial proteins, albeit reduced, suggests that the Spo11 core complex has an ability to bind DSB sites by itself. Whether the same Spo11 core molecule bind both a DSB site and axis site in the tethered loop-axis complexes or different Spo11 core molecules separately bind DSB and axis sites, which are eventually tethered, is difficult to distinguish from our ChlP-seq data. Nonetheless, we infer that initial assembly of each complex at chromosome axes and DSB hotspots via distinct pathways is further strengthened by inter-complex interactions such as Mre11-Mer2, Mre11-Spo11, and Rec114-Rec102/104 ^25, 31, 39, 41^ in the context of tethered loop-axis complexes to efficiently induce DSBs.

### Multi-layered control of chromosomal assembly of the DSB machinery contributes to shaping the DSB landscape in budding yeast

Our correlation analysis with DSB density in variable window sizes in budding yeast provides mechanistic insights into a link between DSB distribution and chromosomal distributions of DSB and axial proteins at chromosomal domain scale, which is distinctly regulated from their local assembly and also controlled via distinct mechanisms at short (≤ 7.5 kb) and long (> 7.5 kb) distances. The estimated average loop length of ∼14 kb ^16^ implicates that the short domains correspond to intra-loop regulation while the long domains include both intra- and inter-loop regulations. We infer that initial local assembly of the full DSB machinery is followed by their redistribution or modulation to generate chromosomal domains 30-40 kb wide ranging multiple axis-loop units where all of them are enriched and DSBs form with higher frequency (**Fig. 6i**). Dependency of Spo11 and Red1 distributions on RMM components and Hop1 suggests that inter-complex interactions that are not required for local assembly of the DSB machinery are critical for this process. This may involve interactions between the DSB machinery in multiple axis-loop units (i.e. inter-loop interactions). Although the precise mechanisms of this redistribution/modulation process remain to be addressed, our study provides multi-layered mechanisms of chromosomal assembly of the DSB machinery from initial local recruitment/assembly to short (intra-loop)- and long (inter-loop)-range distributions, which contributes to shaping the DSB landscape.

### Similarity and distinction of DSB regulation at chromosomal domain scale between budding yeast and mice

For window sizes ≤ 7 kb, all DSB and axis-organizing proteins in mouse spermatocytes analyzed in this study showed an increase in a positive correlation with DSB density, with higher correlations for DSB and axial proteins than cohesin and CTCF. This suggests a conserved mechanism of DSB regulation by the amount of DSB and axis-organizing proteins at short, intra-loop distance between budding yeast and mice. With large windows, in contrast, SYCP3, HORMAD1, and IHO1 showed a decrease in correlations up to ∼100 kb, while MEl4, REC8, RAD21L, and CTCF showed a further increase in correlations. Expanding window sizes to Mb scale given the longer loop lengths in mice revealed a high positive correlation with DSB density for HORMAD1, MEl4, and CTCF, a modest positive correlation for REC8 and RAD21L, and little or rather a weak negative correlation for SYCP3 and IHO1. Thus, only a part of DSB and axial proteins are enriched at DSB-hot chromosomal domains corresponding to sizes of the axis-loop units in mice, in sharp contrast to enrichment of all DSB and axial proteins at those domains in budding yeast. This distinction could be partly explained by distinct mechanisms of the assembly of axial proteins and RMM/RMl components on meiotic chromosomes between budding yeast and mice. While Hop1 is lost from chromosomes in the absence of Red1 ^15^, HORMAD1 can localize to chromosome axes in the absence of SYCP2 and SYCP3, likely via physical interaction with cohesin complexes ^63, 64^. Mer2 forms punctate immunostaining foci ^31, 65^ and controls chromosomal localization of its partner Mei4, but not Mer2 itself, via DDK-dependent phosphorylation, in contrast to IHO1 that requires DDK-dependent phosphorylation for its axis localization as linear immunostaining signals ^66^. High variability of local binding patterns of DSB and axial proteins at chromosome axes in mice, in contrast to their similar, but not identical, enrichment at where cohesin Rec8 preferentially localizes in budding yeast, also supports the idea that mechanisms of the chromosomal assembly of DSB and axial proteins are distinct between the two species.

Dependency of MEl4 distribution at chromosomal domain scale, but not its local enrichment, on SPO11 that forms a distinct complex from MEl4 ^20^ suggests that inter-complex interactions also play critical roles in the regulation of chromosomal domain-scale distributions of DSB proteins in mice. Furthermore, dependency of HORMAD1 distribution on SPO11 and IHO1 suggests interdependent, rather than simple dependency, relationships between axial and DSB proteins on their chromosomal distributions.

Although the mechanisms of initial recruitment/assembly of DSB and axial proteins at chromosome axes and DSB hotspots remain to be fully understood in mice, we infer that initial local assembly of the full DSB machinery in mice also undergoes further modulation or redistribution via inter-complex interactions (e.g. SPO11-MEl4 and SPO11-HORMAD1) to generate chromosomal domains 2-3 Mb wide, which also ranges multiple axis-loop units, where a part of them are enriched and DSBs form with higher frequency (**Fig. 6i**).

While correlation values for chromosomal domain scale that correspond to sizes of the axis-loop units reached Pearson’s *r* = ∼0.8 at 40-kb windows in budding yeast, those in mouse spermatocyte were Pearson’s *r* = ∼0.6 at 3-Mb windows. This suggests that chromosomal domain-scale distribution of the DSB machinery is more critical for controlling DSB distribution in budding yeast while other factors such as their local enrichment and/or local chromatin structures contribute more to DSB regulation in mice. Further comprehensive distribution analysis of DSB proteins including SPO11 and TOPOVlBL in wild type and mutants will provide more insights into the multi-layered mechanisms of DSB regulation in mammalian meiocytes.

### Binding of axial proteins to DSB sites is linked to DSB repair

Our analysis revealed Red1-FLAG binding to both DSB hotspots and chromosome axes, especially at later timepoints when pro-crossover factors ZMMs (Zip2, Zip3, Zip4, Spo16, Msh4, Msh5 and Mer3) bind chromosomes ^67–71^. A high correlation of Red1-FLAG enrichment at DSB hotspots with Mer3, but not Spo11, indicates distinct behavior of Red1 from DSB proteins and coordinated action of Red1 with pro-crossover factors at crossover-designated sites for DSB repair. The reason why previous ChlP analysis using Red1 antibody and N-terminally V5-tagged Red1 ^15, 17^ did not detect Red1 binding at DSB hotspots is unknown. One possible explanation is that the C-terminal region of Red1 is in closer proximity to DSB sites and our ChlP analysis for C-terminally FLAG-tagged Red1 allowed efficient detection of its binding to DSB sites. Another possibility is that previous studies, which performed ChlP at 3 hr ^15^ or 4 hr ^17^ after meiotic induction, detected Red1 distribution before its binding to crossover-designated sites was evident due to a slight difference of the progression of meiotic time course from this study.

A recent study in mouse spermatocytes revealed SYCP3 binding at not only the center of DSB sites but also resected DSB ends ^59^. Whether or not SYCP3 binds crossover-designated sites like Red1-FLAG remain unsolved because ChlP for ZMM proteins in mice has not been reported. Moreover, SYCP3 binding to the center of DSB hotspots does not require DSB proteins or DSB activity ^59^, in contrast to DSB-dependent binding of Red1-FLAG to DSB hotspots. Although the mechanisms that regulate binding of Red1-FLAG and SYCP3 to DSB sites seem distinct, these observations suggest that the binding of axial proteins to DSB sites is tightly linked to DSB repair in both yeast and mice.

## Methods

### Yeast strains and culture

Budding yeast *S. cerevisiae* strains were of the SK1 background. All strains used in this study are listed in Table S1.

Synchronous meiotic culture of bidding yeast was performed as described previously ^8, 16^. Briefly, diploid cells were vegetatively grown in YPD medium, precultured in presporulation medium (SPS) and shifted to sporulation medium (SPM) at 30°C to induce meiosis.

### Chromatin immuno-precipitation (ChIP)-sequencing

ChlP was performed as described previously ^16, 56^. 5-10 x 10^8^ cells of meiotic cells were harvested at indicated timepoints after meiotic induction. Cells were fixed with 1% formaldehyde (Sigma) for 10 min at room temperature, and crosslinking was quenched by adding 2.5 M glycine to a final concentration of 125 mM for 5 min. Cells were washed twice with ice-cold TBS (20 mM Tris-HCl pH 7.6, 150 mM NaCl), and the cell pellet was snap-frozen on liquid nitrogen and stored at -80℃. The frozen cell pellet was suspended in 200 µL of lysis 140 buffer (50 mM HEPES-KOH pH 7.5, 140 mM NaCl, 1 mM EDTA, 1% Triton X-100, 0.1% sodium deoxycholate) supplemented with protease inhibitor (Roche, 04693159001) and disrupted using zirconia beads (0.5 mm, YASUl KlKAl) and a Multi-Beads Shocker (Yasui Kikai) with four rounds of shaking at 2,300 rpm for 1 min at 4℃. The lysate was split into two ∼130 µL aliquots and sonicated on the Covaris M220 for 5 min 30 sec at 5-9°C with a setting of Peak Power 50.0, Duty Factor 20.0, and Cycles/Burst 200 (or equivalent settings on the Covaris S220) to obtain genomic DNA fragments averaging 300 bp. Sheared nuclei were combined, and after adding 300 µL of lysis 140 buffer supplemented with protease inhibitor, centrifuged at 15,000 rpm for 15 min at 4℃ to remove insoluble debris. The chromatin-containing supernatant was mixed with 80 µL of Dynabeads protein A bound to 3-6 µL of FLAG or HA antibody (Wako and Covance, respectively) and incubated for 3.5-4.5 hrs on a rotating wheel at 4℃ for immunoprecipitation. Beads were washed twice with 1 mL of cold lysis buffer 140, once with 1 mL of cold lysis 500 buffer (50 mM HEPES-KOH pH 7.5, 500 mM NaCl, 1 mM EDTA, 1% Triton X-100, 0.1% sodium deoxycholate), twice with 1 mL of cold LiCl/detergent (10 mM Tris-HCl pH 8.0, 250 mM LiCl, 1 mM EDTA, 0.5% Nonident P-40, 0.5% sodium deoxycholate), and once with 1 mL of cold TE (10 mM Tris-HCl pH 8.0 and 1 mM EDTA), followed by elution with 100 µL of TE/1% SDS (50 mM Tris-HCl pH 8.0, 10 mM EDTA, 1% SDS) for 15 min at 65℃ with vortexing every 5 min and 150 µL of TE/0.67% SDS (50 mM Tris-HCl pH 8.0, 10 mM EDTA, 0.67% SDS) for 5 min at 65℃. Two elutes were combined and incubated overnight at 65℃ for de-crosslinking. For input, 10 µL of the chromatin-containing lysate was mixed with 90 µL lysis 140 buffer and 150 of µL TE/1.33% SDS (50 mM Tris-HCl pH 8.0, 10 mM EDTA, 1.33% SDS). Protein was digested by adding 8.4 µL of Proteinase-K (Merck) and incubation for 2 h at 50℃. DNA was purified using QlAquick PCR Purification Kit (QlAGEN).

3-10 ng of ChlP and input DNA was further sonicated on the Covaris M220 for 8 min 30 sec at room temperature with a setting of Peak Power 75.0, Duty Factor 10.0, and Cycles/Burst 200 (or equivalent settings on the Covaris S220) to obtain DNA fragments averaging 50 bp. Multiplexed libraries were prepared with NEBNext ChlP-Seq Library Prep Master Mix Set for Illumina (NEB, E6240S) and NEBNext Multiplex Oligos for Illumina (NEB, E7335S) or NEBNext Ultra Il Library Prep Kit for Illumina (NEB, E7645) and NEBNext Multiplex Oligos for Illumina (NEB, E6440) according to the manufacturer’s instruction. ChlP and input libraries were sequenced on Illumina MiSeq in a 50 bp single-end using MiSeq Reagent Kit v2 (4 samples per run) or NovaSeq6000 in a 150 bp paired-end run. Around 7 million reads were generated per sample.

### ChIP-seq data processing

Single-end 50 bp reads were filtered, end trimmed, followed by removal of the reads containing tag sequences as described previously ^16^. Paired-end 150 bp reads were processed using fastp ^72^. Processed reads were mapped to the *S. cerevisiae* reference genome (sacCer2) using bowtie2 ^73^ with default settings. DNA enrichment (ChlP/input) was calculated in 10-bp non-overlapping bins with 600-bp smoothing windows using Deeptools ^74^. Heatmaps were generated using Deeptools.

ChlP density on each chromosome was calculated by dividing total ChlP signals with total length of mapped bins. Ribosomal DNA (rDNA) cluster (coordinates 450,000-490,000 on chromosome Xll) was excluded.

### Peak classification for ChIP-seq in mice

For peak detection, MACS3 was run for published ChlP-seq data with the following parameters: REC8, -c 25; RAD21L, -c 12; CTCF, -c 15 ^60^; CTCF, -0.5 ^59^. Potential axis sites were defined as follows: CTCF-only sites, CTCF peaks of either leptonema/zygonema ^59^ or pachynema/diplonema ^60^ that overlap with neither REC8 nor RAD21L peaks; CTCF + cohesin, CTCF peaks of either leptonema/zygonema or pachynema/diplonema that overlap with either REC8 or RAD21L peaks; Cohesin-only sites, either REC8 or RAD21L peaks that do not overlap with CTCF peaks of either leptonema/zygonema or pachynema/diplonema. Peaks that overlap with ±2 kb of the center of DSB hotspots defined by SPO11-oligo mapping ^75^ were excluded from those three classes.

## Statistical analysis

Statistical analyses were performed using Graphpad Prism software version 9.5.1 and R version 4.2.2 (https://www.r-project.org). Bars in figures and values in manuscripts indicate means otherwise mentioned. Statistic parameters and tests are described in the figures and/or figure legends.

## Acknowledgments

We thank members of the Ohta lab and Shinohara lab, especially Drs. H. Sasanuma, K. Kugou, and A. H. Oda. We also thank Drs. S. Keeney and B. de Massy for discussion. This work was supported by JSPS KAKENHI Grant Numbers; 24K02068 to K.O. and 24K09545 to M.l. S.M. was supported by Japanese government scholarship and Institute for Protein Research, The University of Osaka. A.S. was supported by grants from Takeda Science Foundation, The Uehara Memorial Foundation, Daiichi Sankyo Foundation of Life Science, The NOVARTlS Foundation Japan for the Promotion of Science, and The Institute for Fermentation, Osaka.

## Author contributions

M.I. and K.O. conceived and designed the experiments. M.I. and S.M. performed ChIP-seq experiments. M.I. and A.S. analyzed the data. M.I. wrote the manuscript with inputs and edits from all authors.

## Competing interest

The authors declare no competing interests.

## Notes

### Competing Interest Statement

The authors have declared no competing interest.

