## Supplementary figures and table for "Multi-layered control of chromosomal assembly of the meiotic DNA break machinery"

Ito et al, Figure S1

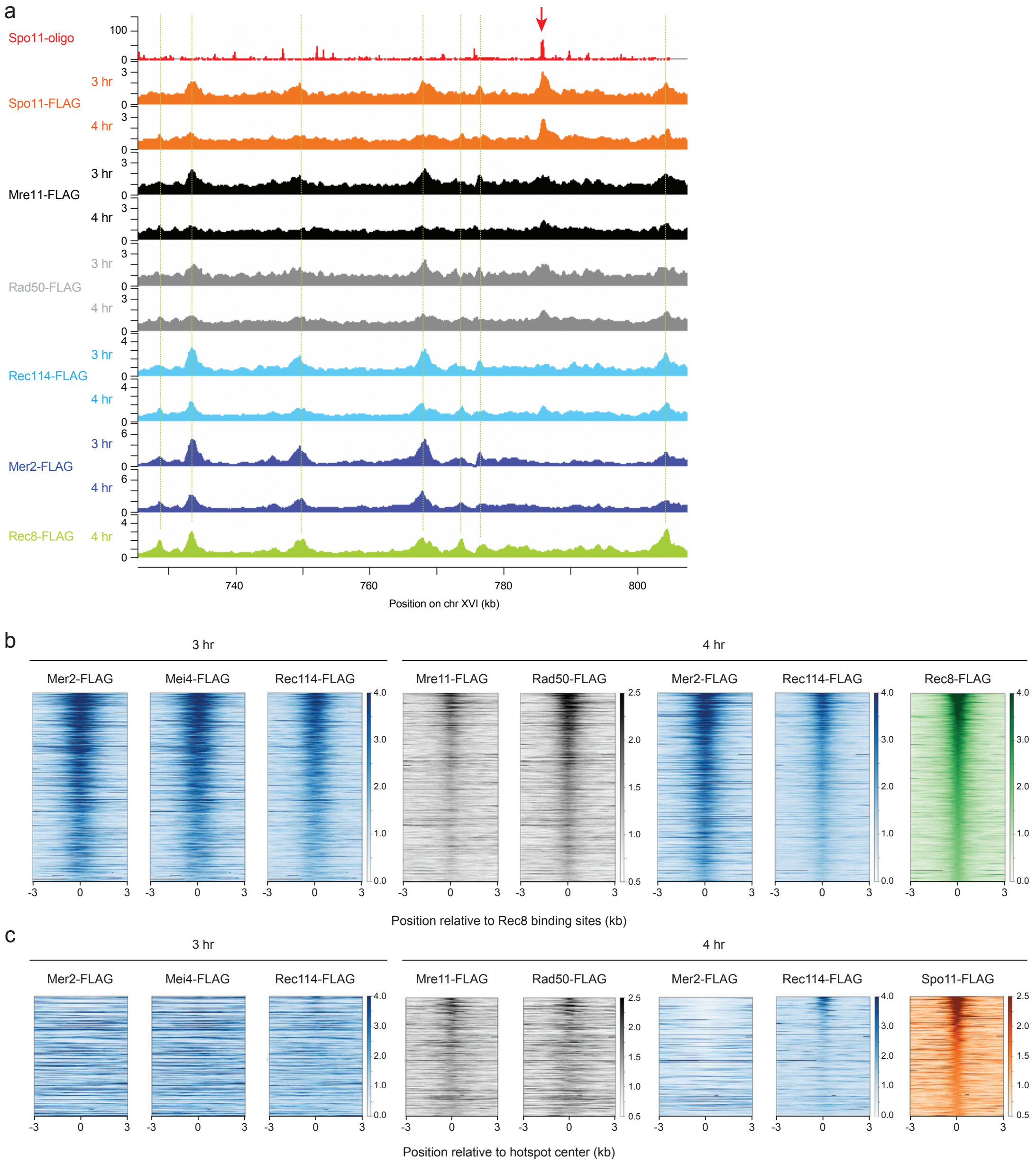


**Fig. S1. Localization of MRX and RMM components to chromosome axes and DSB hotspots.**

(a) ChIP signals of MRX and RMM components in wild type at 3 hr and 4 hr after meiotic induction at the same chromosomal region shown in Fig. 1a. Spo11-oligo sequencing data and ChIP signals of Spo11-FLAG and Rec8-FLAG are reproduced from Fig. 1a to aid comparison. A red arrow and green dashed lines indicate a DSB hotspot with prominent Spo11 enrichment and Rec8 binding sites, respectively.

(b) Heatmaps of ChIP signals of MRX and RMM components in wild type at indicated timepoints after meiotic induction around chromosome axes. The top 540 sites with highest Rec8 enrichment in wild type at 3 hr and 4 hr among 724 Rec8 binding sites identified in our previous study [16] are ordered by Rec8 enrichment. ChIP signals are centered relative to Rec8 binding sites.

(c) Heatmaps of ChIP signals of MRX and RMM components in wild type at indicated timepoints after meiotic induction around DSB hotspots. The top 360 sites with highest Spo11 enrichment in wild type at 3 hr and 4 hr among 3,600 DSB hotspots identified in a previous study [50] are ordered by Spo11 enrichment. ChIP signals are centered relative to hotspot centers.

Ito et al, Figure S2

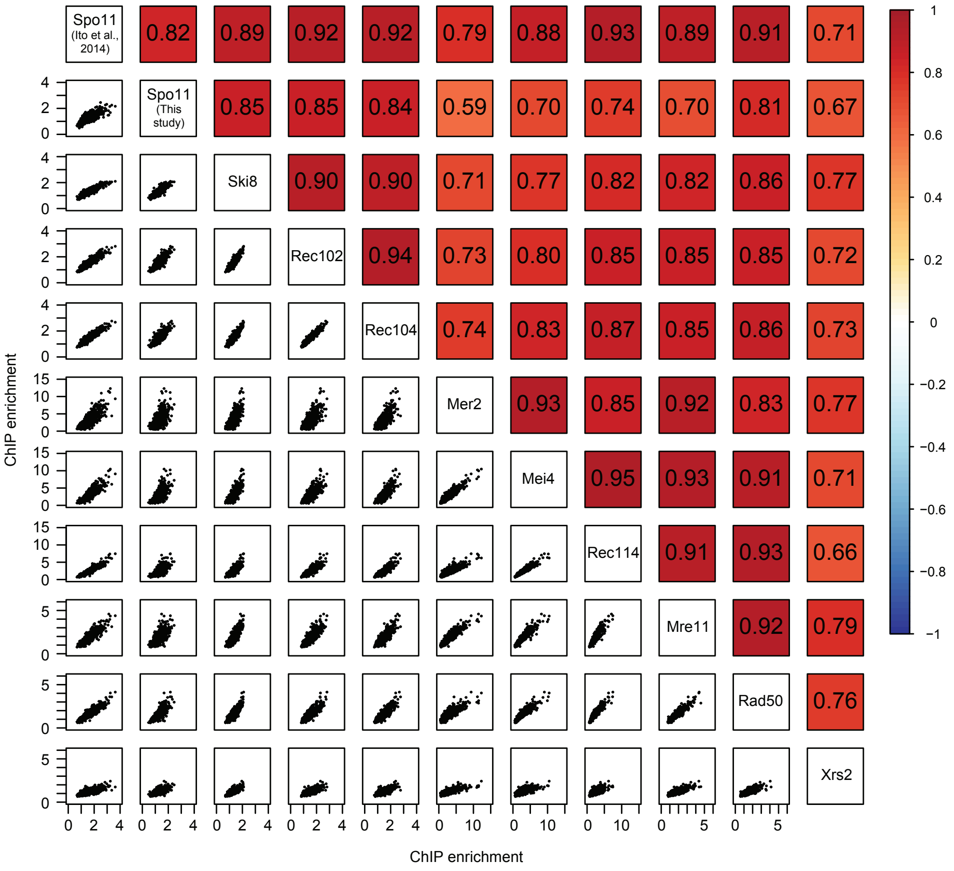


**Fig. S2. Similar distributions of DSB proteins at chromosome axes.**

Correlations of ChIP signals at chromosome axis sites between DSB proteins in wild type at 3 hr after meiotic induction. Each dot indicates one of the 724 Rec8 binding sites identified in our previous study [16]. Pearson's *r* values are also indicated.

Ito et al, Figure S3

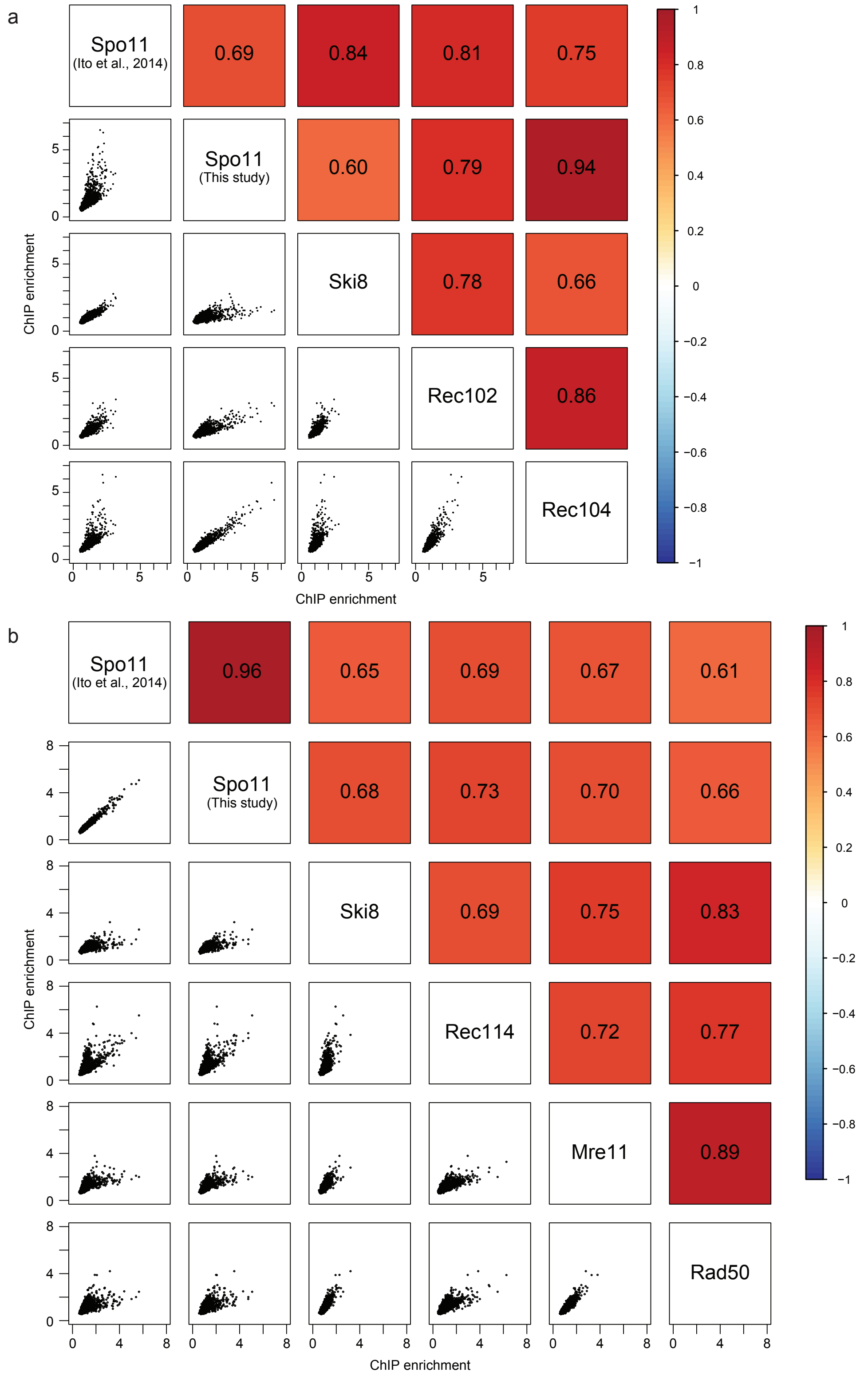


**Fig. S3. Similar distributions of DSB proteins at DSB hotspots.**

(a) Correlations of ChIP signals at DSB hotspots between Spo11 core components in wild type at 3 hr after meiotic induction from our previous study [16] and this study. Each dot indicates one of the 3,600 DSB hotspots identified in a previous study [50]. Pearson's *r* values are also indicated.

(b) Correlations of ChIP signals at DSB hotspots between indicated DSB proteins in wild type at 4 hr after meiotic induction from our previous study [16] and this study, displayed as in (a).

Ito et al, Figure S4

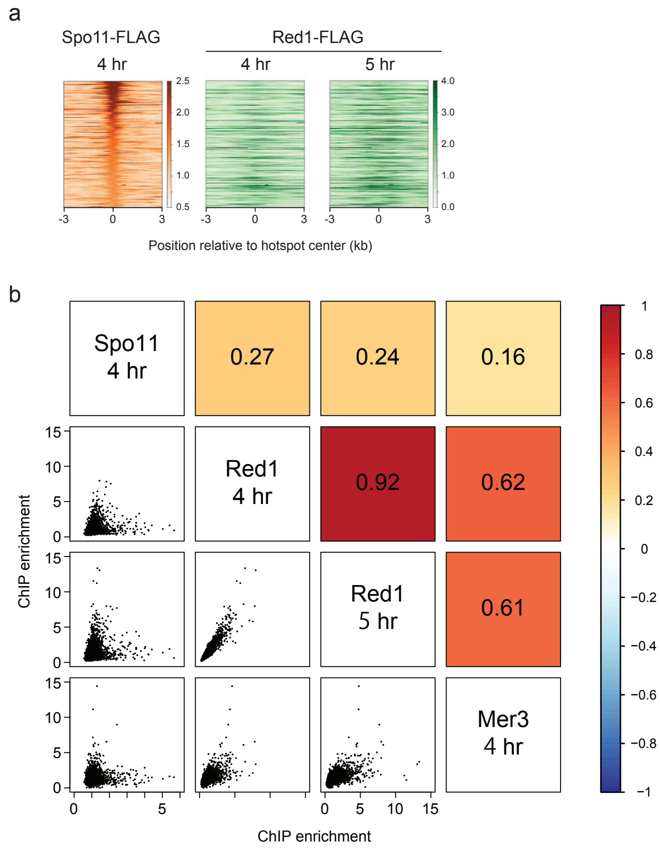


**Fig. S4. Similar distribution of Red1 to a pro-crossover factor Mer3, but not Spo11, at DSB hotspots.**

(a) Heatmaps of ChIP signals of Red1-FLAG in wild type at 4 hr and 5 hr after meiotic induction around DSB hotspots. The top 360 sites with highest Spo11 enrichment in wild type at 4 hr among 3,600 DSB hotspots identified in a previous study [50] are ordered by Spo11 enrichment. ChIP signals are centered relative to hotspot centers.

(b) Correlations of ChIP signals at DSB hotspots between Spo11-FLAG, Red1-

FLAG, and Mer3-FLAG in wild type at indicated timepoints after meiotic induction from previous studies [16, 58] and this study, displayed as in Supplementary Fig. 3.

Ito et al, Figure S5

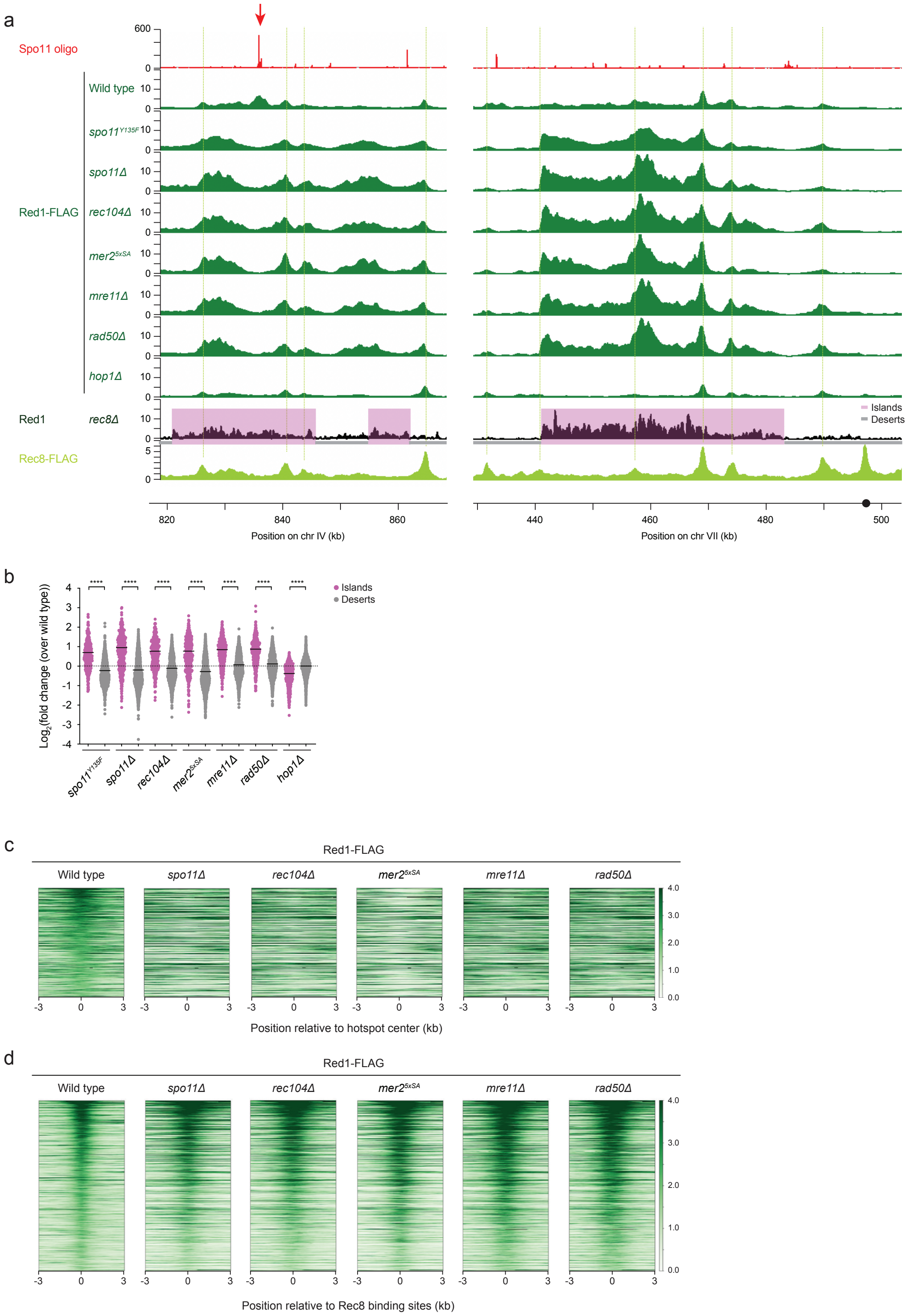


**Fig. S5. Altered Red1 distribution in the absence of DSBs and Hop1.**

(a) ChIP signals of Red1-FLAG in wild type and indicated mutants at 4 hr after meiotic induction at a part of the same chromosomal region shown in Fig.4a and a representative region on chromosome VII. Spo11-oligo sequencing data and ChIP signals of Rec8-FLAG in wild type at 4 hr and Red1 in the *rec8Δ* mutant at 3 hr after meiotic induction from published studies [15, 16, 50] are also shown. A red arrow and green dashed lines indicate a DSB hotspot with prominent Red1 enrichment in wild type and Rec8 binding sites, respectively. Chromosomal regions highlighted in Magenta are previously defined Rec8-independent Red1 binding sites (islands) [15, 18] ; gray lines indicate other regions (deserts), respectively. A black circle indicates the position of centromere.

(b) Log-transformed fold change in Red1-FLAG ChIP signals in indicated mutants over wild type at 4 hr after meiotic induction in islands (magenta) and deserts (gray). Each dot represents one of non-overlapping 5 kb bins excluding rDNA cluster on chromosome XII. Black bars indicate means. The results of the two-tailed Mann-Whitney *U*-test are indicated in the graph: ****$p \leq 0.0001$.

(c, d) Heatmaps of ChIP signals of Red1-FLAG in wild type and indicated mutants at 4 hr after meiotic induction around DSB hotspots (c) and chromosome axes (d), displayed as in Fig. 4b and 4d, respectively.

Ito et al, Figure S6

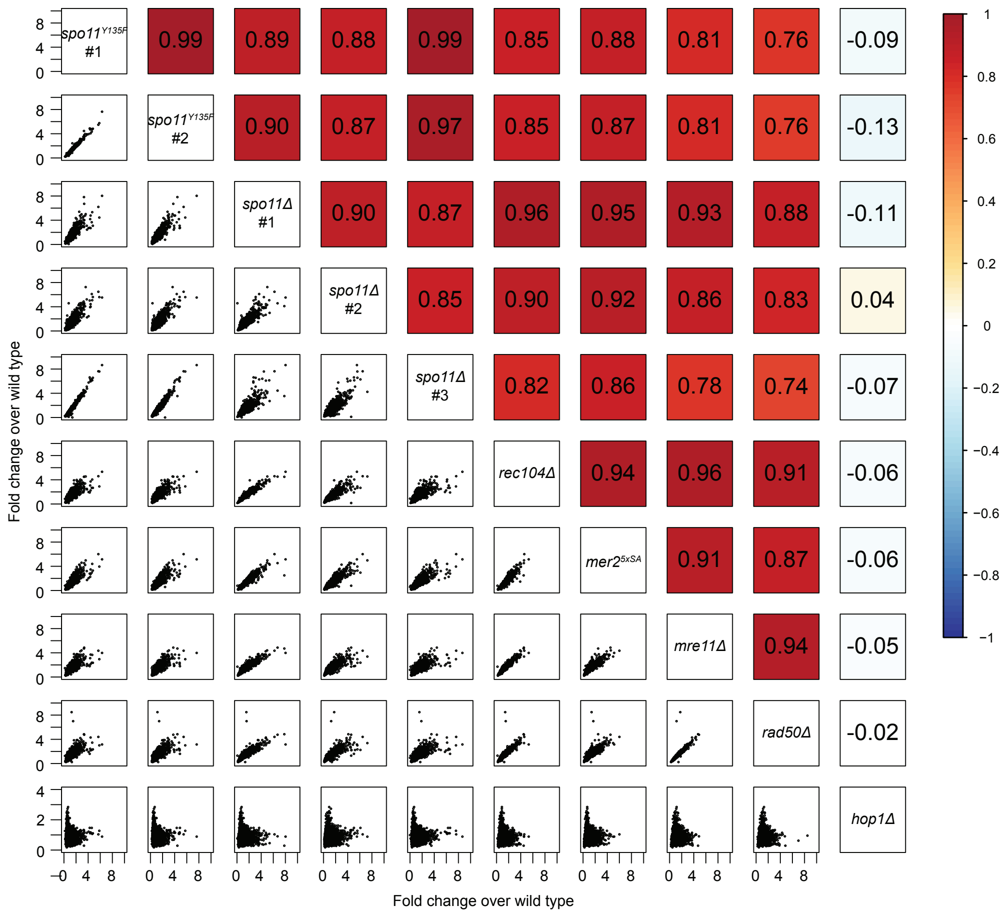


**Fig. S6. Similar changes in Red1 distribution between DSB-defective mutants.**

Correlations of fold-change in Red1-FLAG ChIP signals in indicated mutants (ratio of mutant:wild type) at 4 hr after meiotic induction. Each dot indicates one of 5-kb non-overlapping bins in the entire genome excluding rDNA cluster on chromosome XII. Pearson's *r* values are also indicated.

Ito et al, Figure S7

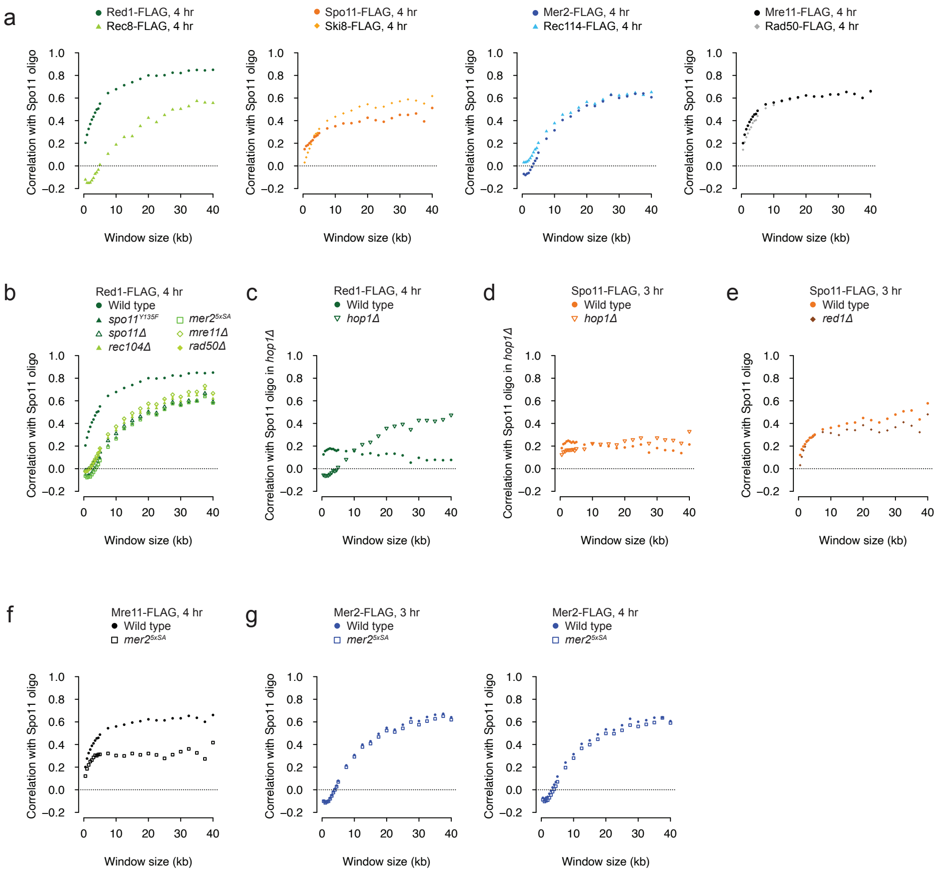


**Fig. S7. Distinct patterns of correlations between ChIP density of DSB and axis-organizing proteins and DSB frequency at chromosomal domain scale in wild type and mutants.**

(a) Scale-dependent correlations of DSB frequency with ChIP density of indicated proteins in wild type at 4 hr after meiotic induction, displayed as in Fig. 5a.

(b) Scale-dependent correlations of DSB frequency with ChIP density of Red1-FLAG in wild type and indicated mutants at 4 hr after meiotic induction, displayed as in Fig. 5a.

(c, d) Correlations of DSB frequency with ChIP density of Red1-FLAG (c) and Spo11-FLAG (d) in wild type and *hop1Δ* mutants at indicated timepoints after meiotic induction, displayed as in Fig. 5a but compared with Spo11-oligo density in *hop1Δ* mutants [10].

(e-g) Scale-dependent correlations of DSB frequency with ChIP density of Spo11-FLAG (e), Mre11-FLAG (f), and Mer2-FLAG (g) in wild type and indicated mutants at indicated timepoints after meiotic induction, displayed as in Fig. 5a.

Ito et al, Figure S8

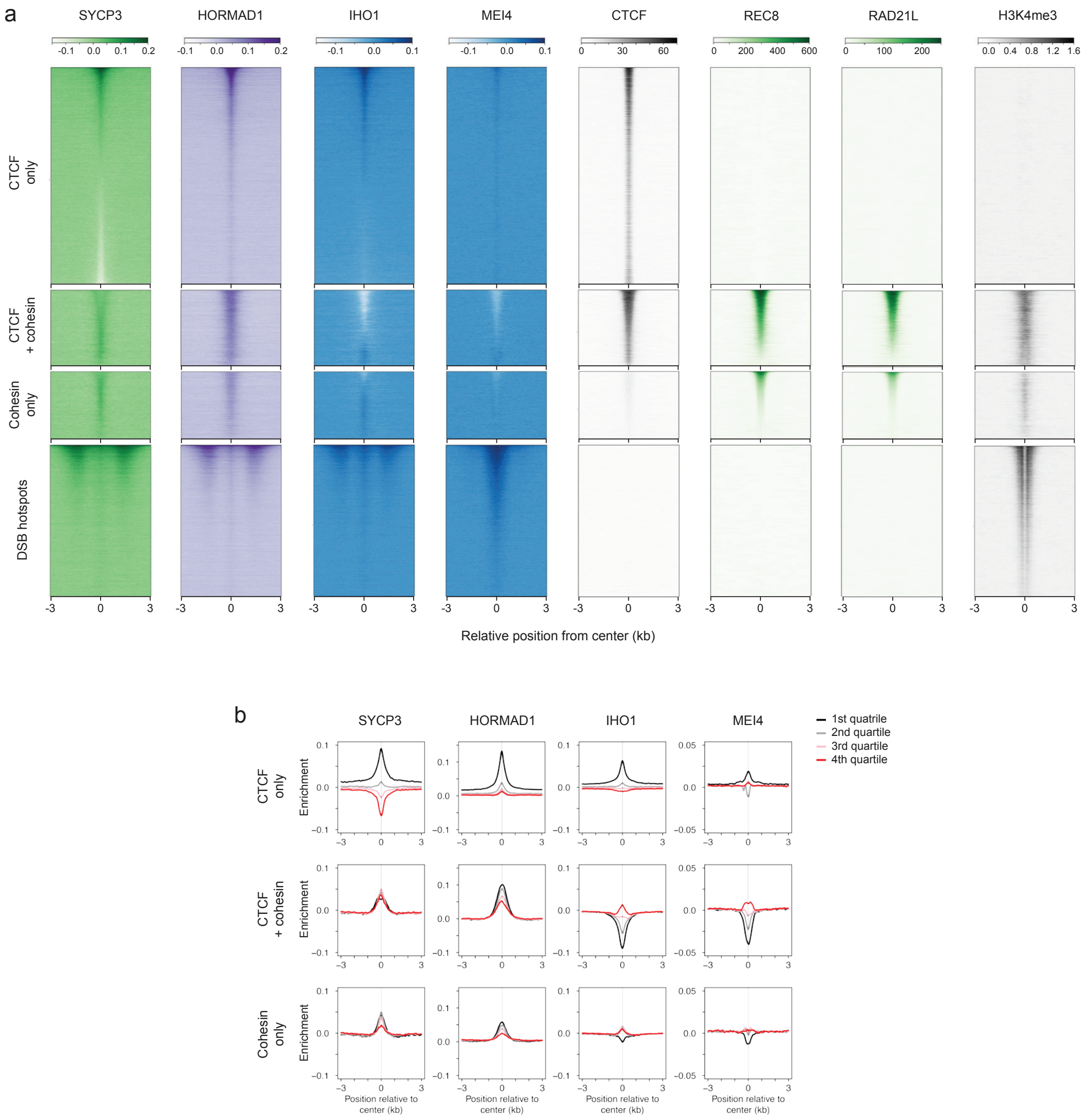


**Fig. S8. Distinct patterns of local distribution of DSB and axis-organizing proteins at chromosome axes and DSB hotspots in mouse spermatocytes.** (a) Heatmaps of ChIP signals of indicated DSB and axis-organizing proteins and histone modification during meiotic prophase I in wild-type mouse spermatocytes from published studies [59, 60, 79]. 19,939 CTCF-only sites, 7,132 CTCF + cohesin sites, 6,379 cohesin-only sites, and 13,944 DSB hotspots identified in a previous study [75] are ordered by SYCP3, REC8, REC8 enrichment, and SPO11-oligo counts, respectively. ChIP signals are centered relative to indicated sites.

(b) Metaplots of ChIP signals of indicated proteins in wild type around chromosome axis sites. Smoothed ChIP signals around ±3 kb of CTCF-only, CTCF + cohesin, and cohesin only sites used for heatmap representation in (a) are shown by dividing the sites in each category into four based on the orders in heatmaps in (a).

Ito et al, Figure S9

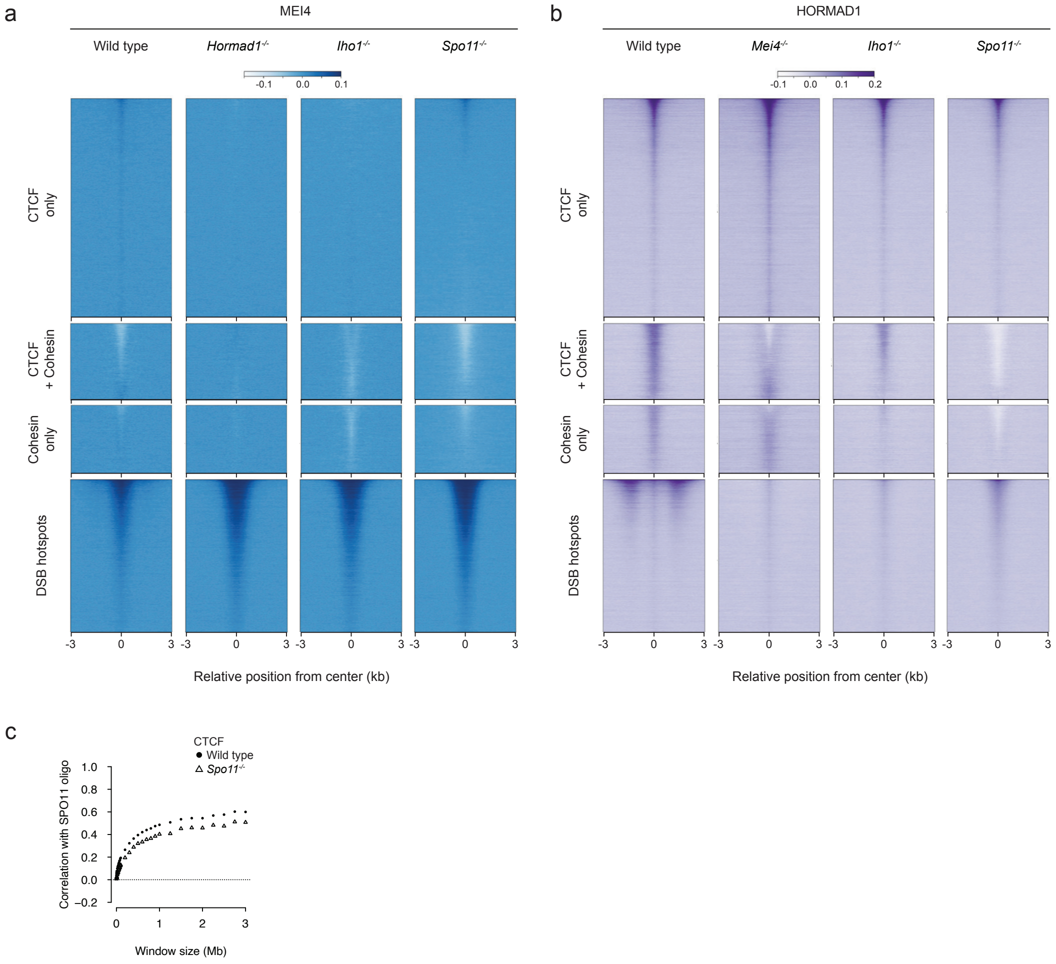


**Fig. S9. Distinct patterns of chromosomal distribution of DSB and axis-organizing proteins in DSB-defective mutants in mouse spermatocytes.**

(a, b) Heatmaps of ChIP signals of MEI4 (a) and HORMAD1 (b) in wild type and indicated mutant spermatocytes from a published study [59], displayed as in Supplementary Fig. 8a.

(c) Scale-dependent correlations of DSB frequency with CTCF ChIP density at chromosomal domain scale in wild-type and *Spo11*$^{-/-}$ mouse spermatocytes, as displayed in Fig. 6b.

**Table S1. Yeast strains used in this study.**

All strains are isogenic to MJL1720 (*Mat**a**/α ura3/" lys2/" ho::LYS2/" leu2Δ/" arg4-bgl/arg4-nsp cyh2-z/"*) with additional modifications described below.

| Strain | Genotype |
|---|---|
| RKD1311 | *SPO11-6His-3FLAG-loxP-KanMX-loxP/"* |
| RKD1313 | *MRE11-6His-3FLAG-loxP-KanMX-loxP/"* |
| YKT190 | *RED1-6His-3FLAG-loxP-KanMX-loxP/"* |
| YMI3 | *REC8-6His-3FLAG-loxP-KanMX-loxP/"* |
| YMI88 | *RED1-6His-3FLAG-loxP-KanMX-loxP/" rec104Δ::CgLEU2/"* |
| YMI90 | *RED1-6His-3FLAG-loxP-KanMX-loxP/" mre11Δ::CgLEU2/"* |
| YMI91 | *RED1-6His-3FLAG-loxP-KanMX-loxP/" rad50Δ::CgLEU2/"* |
| YMI115 | *SPO11-6His-3FLAG-loxP-KanMX-loxP/" hop1Δ::CgLEU2/"* |
| YMI118 | *RED1-6His-3FLAG-loxP-KanMX-loxP/" hop1Δ::CgLEU2/"* |
| YMI132 | *SPO11-6His-3FLAG-loxP-KanMX-loxP/" red1Δ::CgURA3/"* |
| YMI135 | *RED1-6His-3FLAG-loxP-KanMX-loxP/" spo11Δ::URA3/"* |
| YMI319 | *MER2-6His-3FLAG-loxP-KanMX-loxP/" XRS2-6His-3HA-loxP-hph-loxP/"* |
| YMI320 | *mer2-S11,15,19,22,29A-6His-3FLAG-loxP-KanMX-loxP/" XRS2-6His-3HA-loxP-hph-loxP/"* |

| | |
|---|---|
| YMI321 | *RED1-6His-3FLAG-loxP-KanMX-loxP/" mer2-S11,15,19,22,29A/"* |
| YMI324 | *MEI4-6His-3FLAG-loxP-KanMX-loxP/" mer2-S11,15,19,22,29A/"* |
| YMI325 | *MEI4-6His-3FLAG-loxP-KanMX-loxP/"* |
| YMI339 | *MRE11-6His-3FLAG-loxP-KanMX-loxP/" mer2-S11,15,19,22,29A/"* |
| YMI487 | *REC114-6His-3FLAG-loxP-KanMX-loxP/" mer2-S11,15,19,22,29A/"* |
| SMY836/838 | *RED1-6His-3FLAG-loxP-KanMX-loxP/" spo11-Y135F-hph/"* |
| SMY837/839 | *RED1-6His-3FLAG-loxP-KanMX-loxP/" spo11-Y135F-hph/"* |
| YHS16/YHS17 | *REC102-6His-3FLAG-loxP-KanMX-loxP/"* |
| YHS20/YHS21 | *RAD50-6His-3FLAG-loxP-KanMX-loxP/"* |
| YKT34/YHS30 | *REC104-6His-3FLAG-loxP-KanMX-loxP/"* |
| YHS37/YHS38 | *SKI8-6His-3FLAG-loxP-KanMX-loxP/"* |
| YHS41/YHS42 | *REC114-6His-3FLAG-loxP-KanMX-loxP/"* |
| YHS114/YHS115 | *MRE11-6His-3FLAG-loxP-KanMX-loxP/" spo11Δ::URA3/"* |
| YHS471/YHS472 | *SPO11-6His-3FLAG-loxP-KanMX-loxP/" rec102Δ::URA3/"* |
| YHS534/YHS552 | *SPO11-6His-3FLAG-loxP-KanMX-loxP/"* |
| | *mer2Δ::6His-3FLAG-loxP-KanMX-loxP"* |
| YHS536/YHS538 | *SPO11-6His-3FLAG-loxP-KanMX-loxP/"* |
| | *rec114Δ::6His-3FLAG-loxP-KanMX-loxP"* |
